# Phylofactorization: a graph-partitioning algorithm to identify phylogenetic scales of ecological data

**DOI:** 10.1101/235341

**Authors:** Alex D. Washburne, Justin D. Silverman, James T. Morton, Daniel J. Becker, Daniel Crowley, Sayan Mukherjee, Lawrence A. David, Raina K. Plowright

## Abstract

The problem of pattern and scale is a central challenge in ecology. The problem of scale is central to community ecology, where functional ecological groups are aggregated and treated as a unit underlying an ecological pattern, such as aggregation of “nitrogen fixing trees” into a total abundance of a trait underlying ecosystem physiology. With the emergence of massive community ecological datasets, from microbiomes to breeding bird surveys, there is a need to objectively identify the scales of organization pertaining to well-defined patterns in community ecological data.

The phylogeny is a scaffold for identifying key phylogenetic scales associated with macroscopic patterns. Phylofactorization was developed to objectively identify phylogenetic scales underlying patterns in relative abundance data. However, many ecological data, such as presence-absences and counts, are not relative abundances, yet it is still desireable and informative to identify phylogenetic scales underlying a pattern of interest. Here, we generalize phylofactorization beyond relative abundances to a graph-partitioning algorithm for any community ecological data.

Generalizing phylofactorization connects many tools from data analysis to phylogenetically-informe analysis of community ecological data. Two-sample tests identify three phylogenetic factors of mammalian body mass which arose during the K-Pg extinction event, consistent with other analyses of mammalian body mass evolution. Projection of data onto coordinates defined by the phylogeny yield a phylogenetic principal components analysis which refines our understanding of the major sources of variation in the human gut microbiome. These same coordinates allow generalized additive modeling of microbes in Central Park soils and confirm that a large clade of Acidobacteria thrive in neutral soils. Generalized linear and additive modeling of exponential family random variables can be performed by phylogenetically-constrained reduced-rank regression or stepwise factor contrasts. We finish with a discussion of how phylofac-torization produces an ecological species concept with a phylogenetic constraint. All of these tools can be implemented with a new R package available online.

## Introduction

The problem of pattern and scale is a central problem in ecology [27]. Ecological patterns of interest, such as ecosystem physiology, species abundance distributions, epidemics, ecosystem services of animal-associated microbial communities, and more, are often the result of processes that operate at multiple scales. Traditionally, the “scales” of interest are space, time, and levels of ecological organization ranging from individuals to populations to ecosystems. Prediction of spatial variation over different scales, millimeters, meters, or kilometers, requires incorporation of different processes driving patterns observed. Predicting climatic and weather patterns over days, years, or millennia requires different data, processes and models. Predicting the collective behavior of a school of fish requires interfacing individual behavior with interaction networks of those individuals [25] whereas predicting the ability of a forest to act as a carbon sink requires interfacing weather, nutrient cycles, and competition between trees with different traits, such as nitrogen fixation [11]. Understanding emergent infectious diseases requires interfacing processes over scales ranging from animal population dynamics, reservoir epizootiology, and human epidemiology [37]. Ecological theory requires interfacing phenomena across scales believed to be important, and continually updating our beliefs about which scales are important to interface.

For a novel or unfamiliar pattern, such as a change in microbial community composition along environmental gradients, how can one objectively identify the appropriate scales of ecological organization? In macroscopic systems, a researcher will use intuition derived from natural history knowledge to determine scales of interest. Models of how the presumably important natural history traits affect the pattern will be constructed, and the goodness of fit to the pattern of interest will be used as a metric for the successful identification of ecological scales/traits. However, for some patterns, such as the ecosystem physiology of the human microbiome, there is limited natural history knowledge to draw on to assist the decision of the appropriate scales of interest. There is a need for rules, algorithms and laws for the simplification, aggregation, and scaling of ecological phenomena.

A central feature of biological systems is the existence of a hierarchical assemblage of entities, from genes to species, whose relationships and evolutionary history can be estimated and organized into a hierarchical tree. The estimated phylogeny contains edges along which mutations occur and new traits arise. When the phylogeny correctly captures the evolution of discrete, functional ecological traits underlying a pattern of interest, the phylogeny is a natural scaffold for simplification, aggregation, and scaling in ecological systems. Patterns such as the change of bacterial abundances following antibiotic exposure, whose functional ecological traits of antibiotic resistance are laterally transferred, can still be simplified by constructing a phylogeny of the laterally transferred genes, such as the beta-lactamases[18], as a natural scaffold for defining the entities with different responses to antibiotics.

The phylogeny contains a hierarchy of possible scales for aggregation. Graham et al. [17] develop the term “phylogenetic scale” to refer to the depth of the tree over which we aggregate information from a clade. Functional ecological traits often arise at different depths of the tree. Many ecological phenomena may be driven by traits not properly summarized or aggregated by mowing the phylogeny along a constant depth. Instead, there may be multiple phylogenetic scales, or grains, underlying an ecological pattern of interest. For example, the patterns of vertebrate abundances on land and water are simplified by nested clades: Tetrapods, Cetaceans, Pinnipeds, etc. There is a need for general statistical methods to partition the phylogeny into the grains with significantly different associations or contributions to the ecological pattern. Such a method can objectively identify the phylogenetic scales underlying an ecological pattern of interest.

Phylofactorization [51] was developed to identify the phylogenetic scales in compositional (relative abundance) data by iteratively constructing variables corresponding to edges in the phylogeny and selecting variables which maximize an objective function. The variables used were a common transform from compositional data analysis [1], referred to as the isometric log-ratio transform [10, 9], which contrast the relative abundances of species separated by an edge in the phylogeny. A coordinate in an isometric log-ratio transform aggregates relative abundances within clades by a geometric mean and contrasts clades through log-ratios of the clades’ geometric mean relative abundances. The isometric log-ratio transform also allows the construction of non-overlapping contrasts, thereby reducing an obvious source of dependence in phylogenetic variables. The isometric log-ratio transform is used to identify phylogenetic scales capturing large blocks of variation in relative-abundance data and construct coordinates that correspond to edges along which hypothesized functional ecological traits arose.

However, many ecological data are more appropriately viewed as counts, not compositions. For example, the presence/absence of bird species across continents are best modelled as Bernoulli random variables, not compositional data. In this paper, we extend phylofactorization to broader classes of data types by generalizing the logic of phylofactorization and to a set of three operations: aggregation, contrast, and an objective function defined by the pattern of interest. The nested dependence of clades within clades is avoided by defining phylofactorization as a graph-partitioning algorithm that contrasts species separated by edges and iteratively partitioning the phylogeny along edges that best differentiate species.

After defining phylofactorization as a graph-partitioning algorithm, we illustrate the generality of the algorithm through several examples. First, we show that two-sample tests, such as t-tests and Fisher’s exact test, are natural operations for phylofactorization - they first aggregate data from two groups through means, contrast the aggregates via a difference of means, and have natural objective functions defined by their test-statistics. We illustrate the use of two-sample tests by performing phylofactorization of a dataset of mammalian body mass.

Then, we show how the phylogeny serves as a scaffold for changing variables in biological data through a contrast basis - the same basis used in the isometric log-ratio transform - which can be used to identify the phylogenetic scales providing low-rank, phylogenetically-interpretable representations of a dataset. Defining the contrast basis allows us to introduce a phylogenetic analog of principal components analysis - phylogenetic components analysis - which identifies the dominant, phylogenetic scales capturing variance in a dataset. We perform phylogenetic components analysis on the American Gut microbiome dataset (www.americangut.org) and reveal that some of the dominant clades explaining variation in the American Gut correspond to clades within Bacteroides and Firmicutes, thereby providing finer, phylogenetic resolution of a known, major axis of variation in human gut microbiomes found to be associated with obesity [47], age [31] and more. Another phylogenetic factor of variance in the American Gut is a clade of Gammaproteobacteria strongly associated with IBD, corroborating a recent study’s use of phylofactorization to diagnose patients with IBD [49]. The contrast basis can also be used with regression if the data assumed to be approximately normal, log-normal, logistic-normal or otherwise related to the normal distribution through a monotonic transformation. We illustrate regression-phylofactorization through a generalized additive model analysis of how microbial abundances change across a range of pH, Nitrogen, and Carbon concentrations in soils. The resulting contrast basis and its fitted values from generalized additive modeling yield a low-rank representation of biological big-data and translates to clear biological hypotheses aiming to identify the traits driving observed non-linear patterns of abundance across pH [39].

Datasets comprised of non-Gaussian, exponential family random variables can still be analyzed through regression-phylofactorization. We present four methods for generalized regression-phylofactorization in exponential family data. The first method is to use the contrast basis for constrained, reduced-rank regression to obtain a low-rank approximation of coefficient matrices in multivariate generalized linear models. The second uses a two-level factor, a surrogate variable `phylo` indicating which side of an edge a species is found, to define objective functions based on the deviance or the magnitude of the coefficients for the factor-contrast. The third method aggregates exponential family data within clades to marginally stable distributions within the exponential family, and then performs `phylo` factor contrasts described above. The fourth method is a mix of the first and second, developed to have the accuracy of the second method while reducing the computational costs. The mixed method considers `phylo` factors for only a subset of the best edges obtained from reduced-rank approximation of the coefficient matrix.

We finish with a discussion of the challenges, and opportunities, for future development of phylofactorization, and provide an R package - phylofactor - available at https://github.com/reptalex/phylofactor.

## Phylofactorization

Which vertebrates live on land, and which vertebrates live in the sea (Figure 1a)? Most children have enough natural history knowledge to say “fish live in the sea”, thus correctly identifying one of the most important phylogenetic factors of land/sea associations in vertebrates. The statement “fish live in the sea” can be mathematically formalized by noting that one edge in the vertebrate phylogeny separates “fish” from “non-fish” (Figure 1b). Partitioning the phylogeny along the edge basal to tetrapods can separate vertebrates fairly well by land/sea associations. An algorithm for identifying that edge by land/sea associations alone, without requiring detailed knowledge of macroscopic life and morphological and physiological traits, can correctly identify an edge along which functional ecological traits and life-history traits arose. Controlling for the previously identified edge, one might be able to identify the edges basal to Cetaceans and Pinnipeds, tetrapods which live in the sea (Figure 1b). Three edges can capture most of the variation in land/sea associations across thousands of vertebrate species.

**Figure 1:**
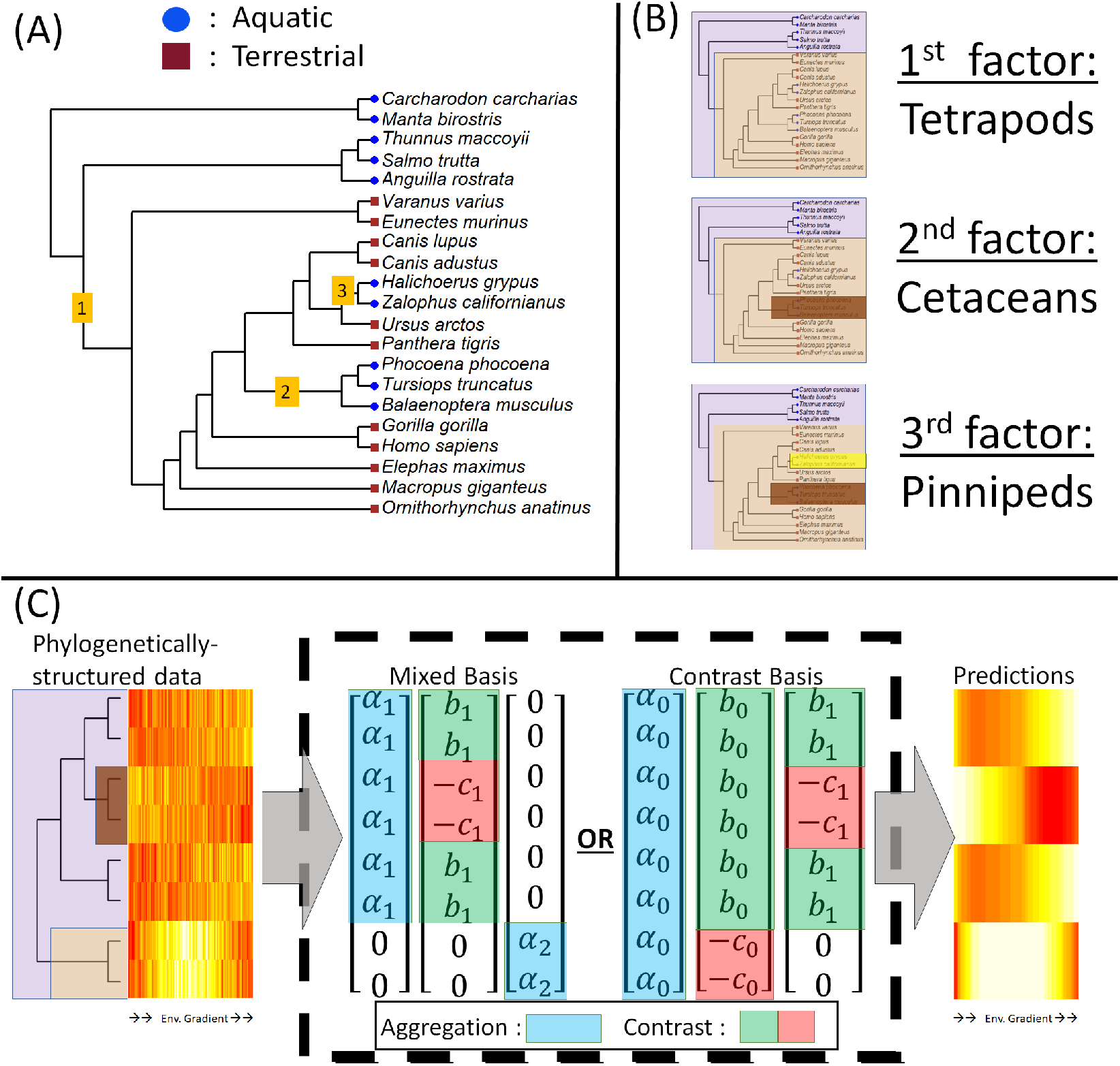
Phylofactorization generalizes the logic of how to simplify phylogenetically-structured datasets. (A) Vertebrate land/water associations can be simplified by partitioning the tree into the edges along which major traits arose. (B) The first phylogenetic factor of vertebrate land/water associations is the edge along which tetrapods arose - an edge along which lungs and limbs evolved that allowed colonization of land. Downstream factors can refine the original partitioning, and include the Cetaceans and Pinnipeds, among other edges along which adaptation to aquatic life arose among tetrapods. (C) Phylogenetic factorization generalizes this same logic for phylogenetically-structured data in which traits might not be known or their evolution easily modeled, including traits like a non-linear relationship between abundance and an environmental gradient. Phylogenetically-structured data can be partitioned through operations of aggregation and contrast. Pure aggregations (blue) sum data within a clade, whereas contrasts (green/red) are differences between two clades. Low-rank, phylogenetically-interpretable predictions of our data can be obtained through a mixed basis of a series of aggregations and contrasts, or a “contrast basis” in which there is a global aggregate partitioned in subsequent contrasts.

Ancestral state reconstruction of habitat association provides a well-known means of making such inferences. However, sometimes the desired traits and ecological patterns of interest are more complicated and their ancestral state reconstruction dubious. For instance, how can we identify the phylogenetic scales of changes in microbial community composition along a pH gradient, allowing possible non-linear associations that could be detected through generalized additive modeling? Answering such a question through ancestral state reconstruction requires conceiving and analyzing an evolutionary model of how the generalized additive models of pH association evolve along a tree. Phylofactorization aims to generalize the phylogenetic logic used for land/sea associations in order to identify phylogenetic scales for more complicated functional traits and ecological patterns, for which an evolutionary model would be dubious. Phylogenetic factorization generalizes the logic of land/sea associations through a graph partitioning algorithm iteratively identifying edges in the phylogeny along which meaningful differences arise (Figure 1c).

### General Algorithm

Phylofactorization requires a set of disjoint phylogenies spanning the set of species considered in the data. The phylogenies are rooted or unrooted graphs with no cycles, containing and connecting the units of interest in our data (the units can be species, genes other evolving units of interest). Phylofactorization can be implemented with disjoint sub-graphs, such as viral phylogenies for which there are not clear common ancestors, and the sub-phylogenies can either be kept separate or joined at a polytomous root. The phylogeny may have an arbitrary number and degree of polytomies.

Let [*x*]_*i,j*_ be the data for species *i* = 1, …, *m* in sample *j* = 1, …, *n*. Let ***x**_R,j_* be the vector of a subset of species, *R*, in sample *j*. Let ***Z*** be the *n* × *p* matrix containing *p* additional meta-data variables for each sample. Let **𝒯** be the phylogenetic tree and let edge *e* partition the phylogeny into disjoint groups *R* and *S*. Phylofactorization requires:

- An aggregation function, 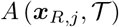 which aggregates any subset, *R*, of species
- A contrast function, 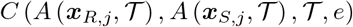 which contrasts the aggregates of two disjoint subsets of species, *R* and *S*, possibly using information from the tree **𝒯** and edge, *e*.
- An objective function, *ω*(*C, **Z***).

With these operations, phylofactorization is defined iteratively as a special case of a graph partitioning algorithm (Figure 2). The steps of phylofactorization are:

1. For each edge, *e*, separating disjoint groups of species *R_e_* and *S_e_* within the sub-tree containing *e*, compute 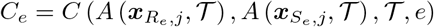
2. compute edge objective *ω_e_* = *ω*(*C_e_, **Z***) for each edge, *e*
3. Select winning edge 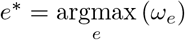
4. Partition the sub-tree containing *e*\* along *e*\*, forming two disjoint subtrees.
5. Repeat 1-5 until a stopping criterion is met.

**Figure 2:**
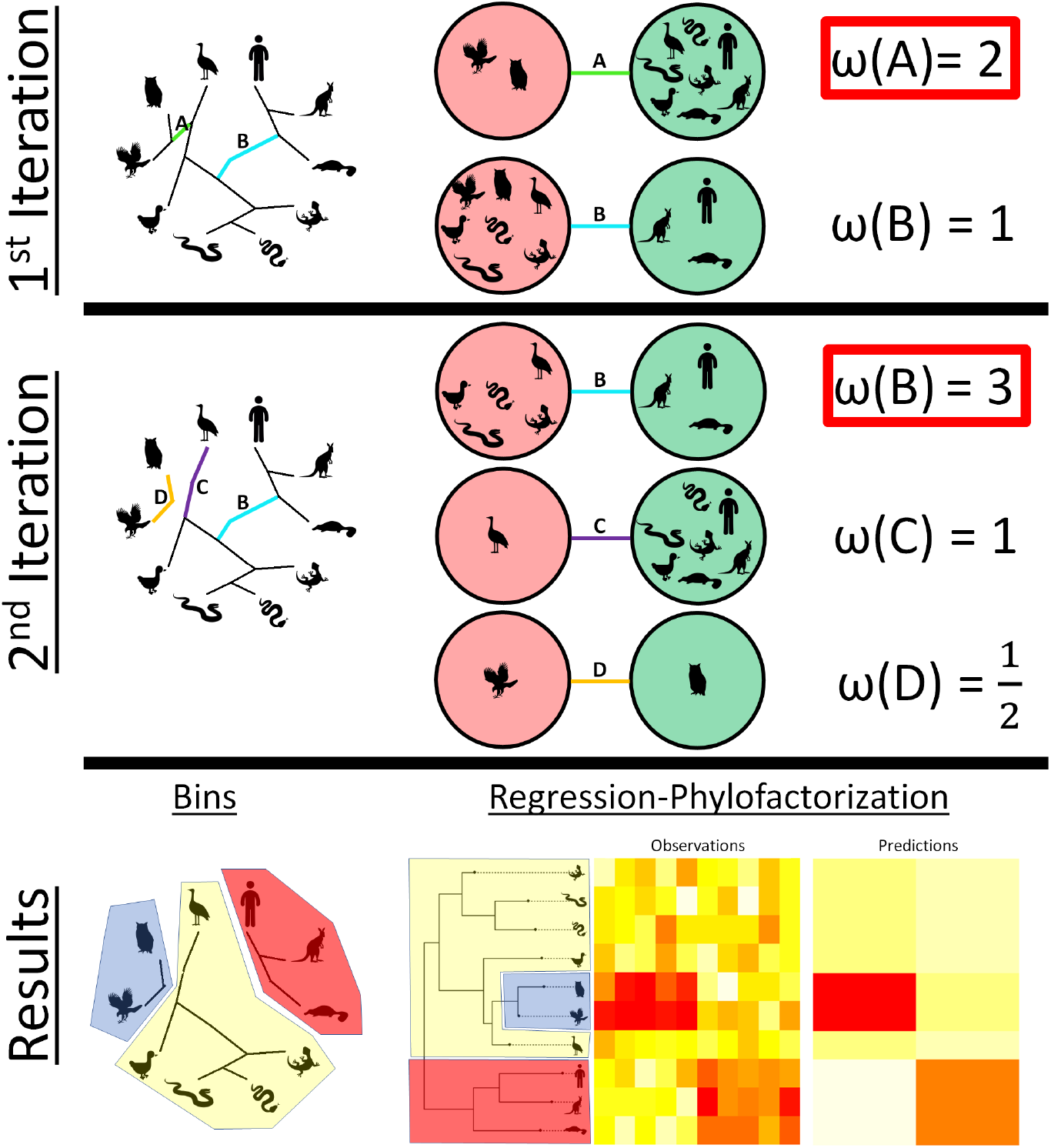
Phylofactorization is a graph partitioning algorithm. An objective function, *ω*, of a contrast of species separated by an edge allows one to iteratively partition the phylogeny along edges maximizing the objective function (1st iteration). After partitioning the phylogeny, the objective functions are re-computed to contrast species in the same sub-tree separated by an edge. Edge B in the first iteration contrasted mammals from non-mammals, but in the second iteration it contrasts mammals from non-mammals, excluding raptors (partitioned in the first iteration). The result of *k* iterations of phylofactorization is a set of *k* + 1 bins of species with similar within-group behavior. A particularly useful case is “regression-phylofactorization”. Regression-phylofactorization is implemented by defining contrasts through the contrast basis (Figure 1c) and defining an objective function through regression on the component scores of each candidate contrast basis element. Regression-phylofactorization is a flexible way to search for clades with similar patterns of association with environmental meta-data while also obtaining low-rank, phylogenetically-interpretable representations of a data matrix.

Unlike more general graph-partitioning algorithms, phylofactorization does not impose a balance constraint - it does not require that the partitions have a similar size or weight. Furthermore, phylofactorization, by working with phylogenies or graphs without cycles is centered around aggregation and contrast as principle operations for defining scales and units of organization. Phylofactorization is limited to contrasts of non-overlapping groups, and the constraint of contrasting aggregates is used to formalize the process of aggregation prior to contrasting groups - such formalization ensures one can subsequently aggregate the bins of species partitioned in phylofactorization according to the method of aggregation by which the bins were discovered to be different. The incorporation of the tree, **𝒯**, in the contrast function encompasses a class of ancestral state reconstruction reconstruction methods. Ancestral state reconstruction with non-overlapping contrasts can be done with time-reversible models of evolution; in this case, phylofactorization contrasts the root ancestral states obtained in which the two nodes adjacent an edge are considered roots of the subtrees separated by an edge.

The edges, *e*\* and their contrasts, *C_e_*, are interchangeably referred to as the “phylogenetic factors” due to their correspondence to hypothesized latent variables (traits) and their ability to construct basis elements that allow matrix factorization [51]. It’s possible to define objective functions through pure aggregation, but we limit our focus to contrast-based phylofactorizations which identify edges along which meaningful differences arose for reasons discussed later in the section on the “contrast basis”.

The result of phylofactorization after *t* iterations is a set of *t* inferences on edges or links of edges. Links of edges occur following a previous partition, when two adjoining edges separate the same two groups in the resultant sub-tree. Partitioning the phylogeny along *t* edges results in *t* + 1 bins of species, referred to as “binned phylogenetic units”. In general, the problem of maximizing some global objective function, 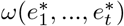, for a set of *t* edges, 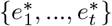, is NP hard [6]. However, stochastic searches of the space of possible partitions, via a stochastic computation of *ω_e_* in step 2 or a weighted draw of *e*\* in step 3, may yield better approximations of a global maximum [32, 20, 23].

Aggregation, contrast, and objective functions are several junctures to define and interpret meaningful quantities and outcomes from data analysis. Explicit decisions about aggregation formalize how a researcher would summarize data from an arbitrary set of species. Explicit decisions about contrast formalize how a researcher differentiates two arbitrary, disjoint groups of species - these common operations form an organizational framework for ecologists studying phylogenetic scales. Aggregation can be done through many operations, including but not limited to addition, multiplication, generalized means, and maximum likelihood estimation of ancestral states under models of trait diffusion away from the focal node. Likewise, examples of contrasts are differences, ratios, various two-sample tests, and more complicated metrics of dissimilarity such as the deviance of a factor contrast in a generalized additive model. Researchers must decide for themselves how best to aggregate information in groups of species, contrast two groups, and decide which group maximizes the objective for a research goal pertaining to a particular ecological pattern. Doing so allows objective, a priori definitions of what makes an informative phylogenetic scale, and the operations chosen are integrated into a broader theoretical framework of phylofactorization.

Below, we develop the generality and illustrate the results from phylofactorization. These examples were run using the R package “phylofactor”, using relevant functions for analyzing and visualizing phylogenies from the R packages ape [36], phangorn [43], phytools [40], and ggtree [53]. Scripts and datasets for every analysis are available in the supplemental materials.

### Example 1: two-sample tests and mammalian body-mass phylofactorization

If the data are a single vector of observations, ***x***, similar to the land/sea associations of vertebrates, phylofactorization can be implemented through standardized tests for differences of means or rate parameters in the two sets of species, *R* and *S*.

To illustrate, we phylofactorize a dataset of mammalian body mass from PanTHERIA [24] and the open tree of life using the R package “rotl” [33]. A single vector of data assumed to be log-normal can be factored based on a two-sample t-test (Figure 3a). In this case, 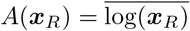 is the arithmetic mean of the log-body-mass; we use the contrast operation

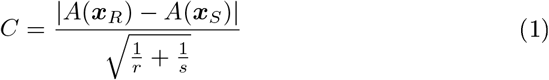

and the objective function *ω_e_* = *C_e_*. Equation (1) defines the test-statistic for a two-sample t-test with the assumption of constant variance. Maximization of the objective function yields edges with the most significant difference in body mass of organisms on different sizes of the tree.

**Figure 3:**
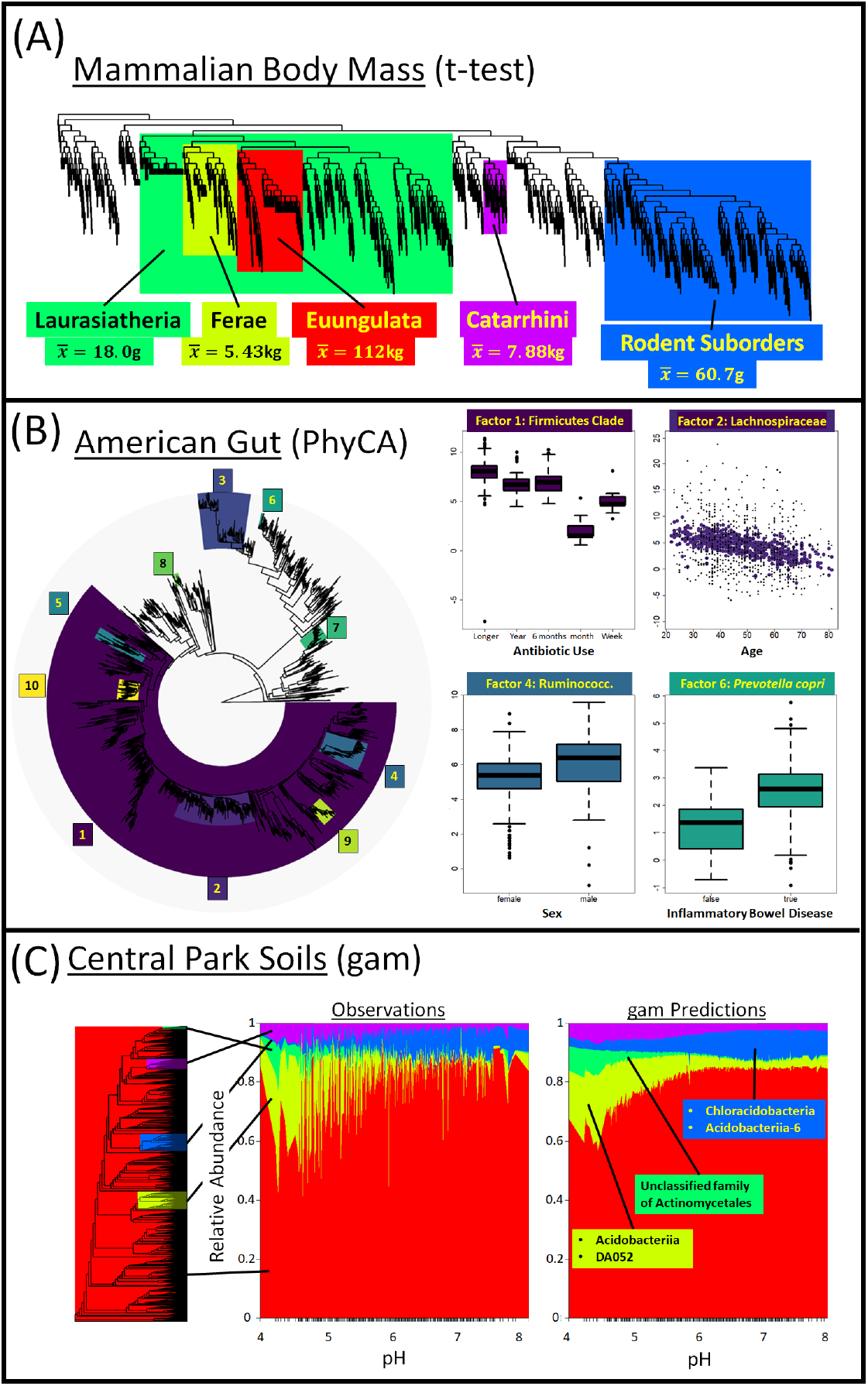
Phylofactorization with contrast basis. (A) The contrast basis defines variables similar to t-statistics, and maximizing the projection of data onto the contrast basis can identify phylogenetic factors. Five iterations of phylofactorization on a dataset of mammalian log-body mass yields five clades with very different body masses. (B) Maximizing the variance of component scores, *y_e_*, of log-relative abundance data produces a “phylogenetic components analysis” (PhyCA) of the American Gut dataset. The most variable clades cover a range of phylogenetic scales. Downstream analysis of component scores tested associations with meta-data - plotted are linear predictors against relevant meta-data; the plot of Lachnospiraceae includes the raw data as black dots. (C) More complicated methods can be used, such as generalized additive modeling with *y_e_*. Using the central park soils dataset, *y_e_* of log-relative abundances, the model *y_e_* ~ *s*(log(Carbon)) + *s*(log(Nitrogen)) + *s*(pH) and the objective of maximizing the explained variance, we obtained the same 4 factors obtained using generalized linear modeling in the original data, including the misnomer group of Chloracidobacteria that don’t thrive in low pH environments. The relative importance of pH in the generalized additive models and exact clades with a high amount of variance explained by pH allows a projection of 3000 species into 5 BPUs for clear visualization of a dominant feature of how soil bacterial communities change along a key environmental gradient.

The first five phylogenetic factors of mammalian body mass in these data are Euungulata, Ferae, Laurasiatheria (excluding Euungulata and Ferae), a clade of rodent sub-orders Myodonta, Anomaluromorpha, and Castorimorpha, and the simian parvorder Catarrhini. Five factors produce six binned phylogenetic units of species with different average body mass (Figure 3a). The most significant phylogenetic partition of mammalian body mass occurs along the edge basal to Euungulata, containing 296 species with significantly larger body mass than other mammals. The second partition corresponds to Ferae, containing 242 species which have body masses larger than other mammals, excluding Euungulata. The third partition corresponds to 864 remaining species in Laurasiatheria, excluding Euungulata and Ferae, which contains Chiroptera, Erinaceomorpha, and Soricomorpha. These mammals have lower body mass than non-Laurasiatherian mammals. The fourth partition identifies three rodent sub-orders comprising 926 species with lower body mass than non-Laurasiatherian mammals. Finally, 106 species comprising the Simian parvorder Catarrhini are factored as having higher body mass than the remaining mammals. These factors are fairly robust: 3000 replicates of stochastic Metropolis-Hasting phylofactorization, drawing edges in proportion to *C^λ^* with *λ* = 6 (producing a 1/4 probability of drawing the most dominant edge) could not improve upon these 5 factors.

The first two phylogenetic factors of mammalian body size partition the mammalian tree at deep edges with ancestors near the K-Pg extinction event, corroborating evidence of ecological release [2, 3] and the exponential growth of maximum body sizes following the K-Pg extinction event [46] for these two dominant clades. The crown group of modern Euungulata are thought to have originated in the late Cretaceous [54] and its representatives may have expanded into previously dinosaur-occupied niches during the rapid evolution of body size in mammals immediately after the K-Pg extinction event at the Cretaceous/Paleogene boundary [45]. Cope’s rule posits that lineages tend to increase in body size over time, and a recent study [4] confirms Cope’s rule and found that mammals have, along all branch lengths in their phylogeny, tended to increase in size. The phylogenetic factors of mammalian body size discovered here illustrate an important feature of phylofactorization: correlated evolution within a clade, such as a consistently high body-size increase among lineages in a clade, can cause the edge basal to a clade to be an important partition for capturing variance in a trait. A more robust phylofactorization may be done through iterative ancestral-state reconstruction of the roots of subtrees partitioned by each edge (where the subtrees are re-rooted at the nodes adjacent the edge), but this unsupervised phylogenetic factorization body masses in 3374 mammals takes 15 seconds on a laptops and yields partitions which simplify the story of mammalian body-mass variation to a set of 5 edges forming 6 binned phylogenetic units.

Two-sample tests can be used for phylogenetic factorization of any vector of trait data. For another example, Bernoulli trait data, such as presence/absence of a trait, can be factored using Fisher’s exact test that there is the same proportion of presences in two groups, *R* and *S*. In this case, the aggregation operation 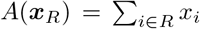 counts the number of successes in group *R*, the contrast operation is the computation of the P-value using Fisher’s exact test with the contingency table

| Successes | Failures | Total |
| --- | --- | --- |
| $A(\mathbf{x}_R)$ | $r - A(\mathbf{x}_r)$ | $r$ |
| $A(\mathbf{x}_S)$ | $s - A(\mathbf{x}_S)$ | $s$ |
| $A(\mathbf{x}_R) + A(\mathbf{x}_S)$ | $r + s - (A(\mathbf{x}_r) + A(\mathbf{x}_S))$ | $r + s$ |

An objective function can be defined as the inverse of the P-value from Fisher’s exact test, 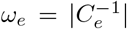. The phylofactorization of vertebrates by land/water association in Figure 1, using an ad-hoc selection of vertebrates for illustration, was performed using Fisher’s exact test, and the factors obtained correspond to Tetrapods, Cetaceans, and Pinnipeds. Unlike the phylofactorization of mammalian body mass, all three factors obtained from phylofactorization of vertebrate land/water association correspond to a set of traits. Tetrapods evolved lungs and limbs which allowed them to live on land. Cetaceans evolved fins and blowholes, and Pinnipeds evolved fins, all traits adaptive to life in the water.

Two-sample tests are used when partitioning a vector of traits and not controlling for additional meta-data such as sampling effort or other confounding effects. Phylofactorization of body mass and land/water associations illustrate two potential evolutionary models under which edges are important: correlated evolution of members of a clade and punctuated equilibria. Edges identified from more complicated methods of phylofactorization may correspond to traits, or they may correspond to directional evolutionary processes shared among members of a clade or their ancestors, such as ecological release or niche partitioning. When the objective function from two-sample tests has a well-defined null distribution, as is the case for the two-sample *t*-test and Fisher’s exact test, the uniformity of the distribution of P-values can used to define a stopping criteria as discussed later (see: “stopping criteria”).

### Example 2: Contrast basis and phylogenetic components analysis

The phylogeny provides a natural scaffold for low-rank, phylogenetically interpretable approximations of the data. As a sphere defines a natural set of coordinates for GPS data, the phylogeny defines a natural set of coordinates that can be used for a variety of data analyses. One example of a natural coordinate in the phylogeny is aggregation: the sum of abundances of species within a clade. Another natural coordinate is a contrast: the differences of abundance between two clades, either sister clades or a monophyletic clade and its complement. Together, these operations allow one to construct natural coordinates for more sophisticated analyses of phylogenetically-structured ecological data.

Phylogenetically-interpretable, low-rank approximations of data can be obtained by constructing basis elements through aggregation and contrast vectors (Figure 1c). An aggregation basis element for a group *Q* = *R* ∪ *S* can be constructed through a vector whose ith element is

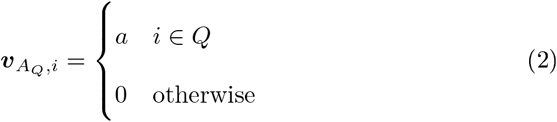

and such aggregation basis elements can be subsequently partitioned with a contrast vector

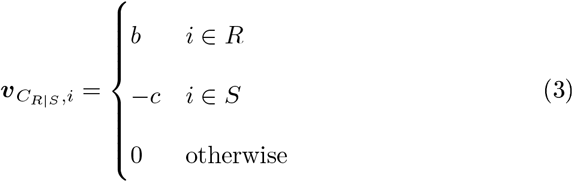

where *b* > 0 and *c* > 0. By meeting the criteria

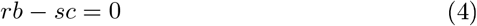

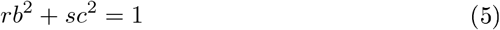

one can ensure that ***v**_A_Q__* and ***v**_C_Q__* are orthogonal and with unit norm. These criteria are satisfied by

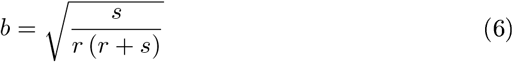

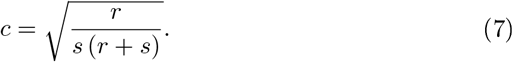

When projecting data from sample *j, **x**_j_*, onto a contrast vector, the aggregation and contrast operations are

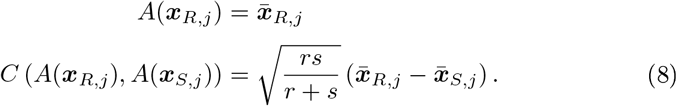

where 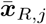 is the sample mean of species in group *R* and sample *j*. Projecting a dataset onto ***v**_C_R|S__* yields coordinates which are a standardized difference of means similar to equation (1). The contrast vector is comprised of two sub-aggregations of opposite sign, one for group *R* and the other for group *S*. By ensuring criterion (4), the groups aggregated within a contrast vector can be subsequently partitioned with additional, orthogonal contrast vectors splitting each group *R* and *S*. Maintaining criterion (5), the aggregation and contrast vectors defined here can be used to construct an orthonormal basis for describing data containing our species, ***x**_j_* ∈ ℝ^*m*^, by defining a set of *q* ≤ *m* orthogonal aggregation vectors corresponding to disjoint sets of species *Q_l_* such that the entire set of aggregations, 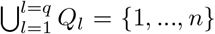, covers the entire set of m species. Then, *m* − *q* contrast vectors partitioning the aggregations and the sub-aggregations within contrast vectors can complete the basis (Figure 1c). Of note is that, as defined in equations (2) and (3), the span of any aggregate and its contrast is equal to the span of the contrasts’ sub-aggregates, i.e. for *R* ∪ *S* = *Q*,

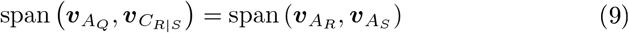

(Figure 1c) and the two natural ways of changing variables with the phylogeny, an aggregate of species and its orthogonal contrast (grouping species and partitioning the group) or two orthogonal aggregates (two disjoint groups of species), are rotations of one-another. Aggregation and contrast vectors translate the notion of phylogenetic scale and group-difřerences into a basis that can be used to analyze community ecological data.

Pure aggregation vectors as defined in equation (2) can be defined a priori based on traits or clades of species thought to be important for the question at hand (e.g. aggregate “terrestrial” and “aquatic” animals), or defined by the data through myriad clustering algorithms or phylofactorization based purely on aggregation by converting steps (1) and (2) in the phylofactorization algorithm into a single step: maximizing an objective function of the aggregate of a clade. A special case occurs when data are compositional [1], in which case the sum of any sample across all species in the community will equal 1 and thus the data are constrained by an aggregation element - the aggregate of all species - which can only be subsequently contrasted. Phylofactorization via contrasts of log-relative abundance data allows one to construct an isometric log-ratio transform, a commonly used and well-behaved transform for the analysis of compositional data [10, 9, 44]. Since the span of an aggregate and its contrast is equal to the span of the contrasts’ two aggregates (equation 9), we simplify construction of the basis by considering, from here on out, only the “contrast basis” in which the an initial aggregate of all species is then partitioned with a series of contrasts.

An orthonormal basis, including one constructed via aggregation and contrast vectors, enables researchers to partition the variance along each of a set of orthogonal directions corresponding to discrete, identifiable features in the phylogeny. Using the phylofactorization algorithm, a dataset ***X*** = [*x*]_*i,j*_ can be summarized by defining the objective function

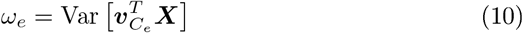

where ***v**_C_e__* is the contrast vector from (3) corresponding to the sets of species, *R* and *S*, split by edge *e*. The objective function in equation (10) yields a phylogenetic decomposition of variance we define as “phylogenetic components analysis” or PhyCA. PhyCA is a constrained version of principal components analysis, allowing researchers to identify the dominant axes of variation, constrained to axes which contrast species separated by an edge.

The variance of component scores, 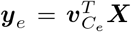, are easiest to understand when the data [*x_i,j_*] are assumed to be standard Gaussian. The component score for sample *j*, ***y**_e,j_*, can be written as

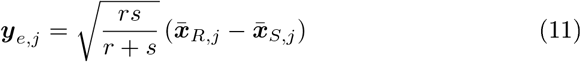

where 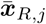 is the sample mean of *x_i,j_* for *i* ∈ *R* and 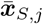 is the sample mean of *x_i.j_* for *i* ∈ *S*. The variance of the component score across all samples *j* = 1, …, *n* is

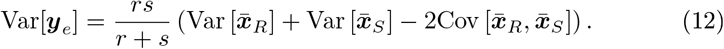

The variance of ***y**_e_* increases through a combination of variances in aggregations of groups *R* and *S* across samples (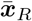 and 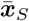, respectively) and a high negative covariance between aggregations for groups *R* and *S* across samples. Species with a negative covariance may be competitively excluding one-another or may be differentiated due to a trait which arose along edge *e* which causes different habitat associations or responses to treatments. Edges extracted from PhyCA are edges along which putative functional ecological traits arose differentiating the species in *R* and *S* in the dataset of interest.

#### Phylogenetic Components of the American Gut

To illustrate, we perform PhyCA to identify 10 factors from a sub-sample of the American Gut dataset and the greengenes phylogeny [8] containing *m* = 1991 species and *n* = 788 samples from human feces (Figure 3b). The American Gut dataset was filtered to only fecal samples with over 50,000 sequence counts and, for those samples, otus with an average of more than one sequence count per sample. After performing PhyCA, each identified resulting component score, ***y**_e*_*, was assessed for a linear association with seven explanatory variables: types_of_plants (a question asking participants how many types of plants they’ve eaten in the past week), age, bmi, alcohol consumption frequency, sex, antibiotic use (ABX), and inflammatory bowel disease (subset_ibd) (Figure 3b). The raw P-values are presented below, but for a reference, the P-value threshold for a 5% family-wise error rate is 7.1 × 10^−4^.

The first factor splits 1229 species of Firmicutes from the remainder of microbes. The component score for the first factor, 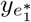, is strongly associated with antibiotic use (P=3.6 × 10^−4^), with dramatic decreases in relative abundance in patients who have taken antibiotics in the past week or month. The second factor identifies 217 species of several genera of Lachnospiraceae, a clade contained within the Firmicutes, strongly associated with age (P=1.2 × 10^−15^) and bmi (P=3.2 × 10^−6^) and alcohol (P=6.4 × 10^−3^). The third factor is a clade of 81 Bacteroides most strongly associated with types_of_plants (P=2 × 10^−9^). By identifying a clade of Bacteroides as a major axis of variation, factors 1 and 3 refine the Firmicutes to Bacteroidetes ratio commonly used to describe variation in the gut microbiome and found associated with obesity and other disease states [28, 7]. It’s been found that the Firmicutes/Bacteroidetes ratio changes with age [31], but the picture from phylofactorization is more nuanced: the large clade of Firmicutes in the first factor does not change with age, but the Lachnospiraceae within that clade decrease strongly with age relative to the remaining Firmicutes, while the Bacteroides show only a moderate decrease with age. The strong decrease with age in Lachnospiraceae is found in a few other clades within the Firmicutes: the 4th factor identified a clade of Firmicutes of the family Ruminococcaceae strongly associated with types of plants (P=3.6 × 10^−5^), sex (P=5.9 × 10^−4^) and decreasing with age (P=9.2 × 10^−4^), and the 5th factor identified a group of Firmicutes of the family Tissierellaceae that decrease strongly with age (P=1.9 × 10^−5^).

The sixth factor is a small group of 5 OTUs of *Prevotella copri* strongly associated with types_of_plants (P=2.8 × 10^−4^) and inflammatory bowel disease (P=2.5 × 10^−3^). Previous studies have found that *Prevotella copri* abundances are correlated with rheumatoid arthritis in people and innoculation of *Prevotella copri* exacerbates colitis in mice. Consequently, *Prevotella copri* is hypothesized to increase inflammation in the mammalian gut [42], and the discovery of *Prevotella copri* as one of the dominant phylogenetic factors of the American Gut, as well as the discovery of its association with IBD, corroborates the hypothesized relationship between *Prevotella copri* and inflammation. Likewise, the seventh factor is a clade of 41 Gammaproteobacteria of the order Enterobacteriales also associated with types_of_plants (P=6.7 × 10^−8^) and weakly associated with inflammatory bowel disease (P=0.022). Gammaproteobacteria were used as biomarkers of Crohn’s disease in a recent study [49] and their associations with IBD in the American Gut project corroborates the possible use of Gammaproteobacterial abundances for detection of IBD from stool samples. Summaries of the models for all factors’ component scores are in the supplemental information.

### Example 3: Compositional, log-normal and Gaussian regression-phylofactorization

The contrast basis can be used to define more complicated objective functions for data assumed to be Gaussian or easily mapped to Gaussian, such as logistic-normal compositional data or log-normal data. Conversion of the data to an assumed-Gaussian form can then allow one to perform least-squares regression using ***y**_e_* as either an independent or dependent variable. Rather than performing PhyCA and subsequent regression, one can choose phylogenetic factors based on their associations with meta-data of interest.

Maximizing the explained variance from regression identifies clades through the product of a high contrast-variance from equation (10) and the percent of explained-variance from regression - such clades can capture large blocks of explained variance in the dataset. Another common objective function is the deviance or *F*-statistic from regression which identifies clades with more predictable responses - such clades can be seen as bioindicators or particularly sensitive clades, even if they are not particularly large or variable clades in the data. Regression-phylofactorization can use the component scores as an independent variable, as was used in the phylofactorization-based classification of Crohn’s disease [49]. For multiple regression, one can use the explanatory power of the entire model, or a more nuanced objective function of a subset of the model. More complicated regression models can be considered, including generalized additive models, regularized regression, and more.

To illustrate the flexibility of regression phylofactorization to identify phylogenetic scales corresponding to nonlinear patterns of abundance-habitat associations, we perform a generalized additive model analysis of the Central Park soils dataset [39] analyzed previously using a generalized linear model. To identify non-linear associations between clades and pH, Carbon and Nitrogen, we perform a generalized additive model of the form

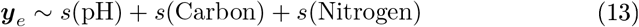

and maximize the explained variance (Figure 3c). The resultant phylofactorizations identifies the same 4 factors as the generalized linear model, but allows nonlinear and multivariate analysis of how community composition changes over environmental meta-data. Combining the high relative-importance of pH with the identified 4 factors, splitting over 3,000 species 5 binned phylogenetic units, allows clear and simple visualization of otherwise complex behavior of how a community of several thousand microbes changes across several hundred soil samples. As with the original analysis, the generalized additive modeling phylofactorization identifies a clade of Acidobacteria - the Chloracidobacteria - which have highest relative abundances in more neutral soils.

### Example 4: Phylofactorization through generalized linear models

Many ecological data are not Gaussian. Presence-absence data or count data with many zeros cannot be easily transformed to yield approximately Gaussian random variables. Data assumed to be exponential family random variables can be analyzed with regression-phylofactorization by adapting concepts in generalized linear models.

We present four options for phylofactorization through generalized linear models. These options correspond to the contrast basis, either explicitly using the contrast basis to approximate the coefficient matrix in multivariate generalized linear models, or implicitly using a form of the contrast basis in the likelihood function when performing shared-coefficient or factor-contrasts in generalized linear modeling.

#### Coefficient Contrast

The first method, related to reduced rank regression for vector generalized linear models [52], uses the contrast basis to provide a reduced-rank approximation of the coefficient matrix from multivariate generalized linear models. Multivariate (vector) generalized linear models assume the data ***X*** are drawn from an exponential family distribution with canonical parameters for each species, ***η*** ∈ ℝ^*m*^, related to the meta-data ***Z*** through a linear model

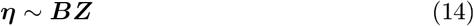

where ***B*** ∈ ℝ^*m*×*p*^ is the coefficient matrix and ***Z*** ∈ ℝ^*p*×*n*^ is the matrix of metadata. Instead of using *m* × *p* coefficients, one can represent the coefficient matrix ***B*** through contrast basis elements and their component scores

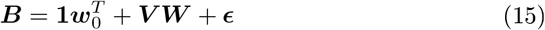

where **1** ∈ ℝ^*m*^ is the one vector, ***w***_0_ ∈ ℝ^*p*^ contains the sum of the regression coefficients for each of the *p* predictors, ***V*** ∈ ℝ^*m*×*t*^ is a matrix whose columns are contrast basis elements obtained from *t* iterations of phylofactorization and ***W*** ∈ ℝ^*t*×*p*^ is a matrix whose rows are the component scores for each contrast basis element. If one is interested in partitioning species based on a subset, P, of the explanatory variables, one can implement equation (15) for the matrix ***B***_P_ containing only the partitioning variables for phylofactorization.

To put multiple independent meta-data from multiple species on the same scale, it’s important to standardize the coefficients *β_i,j_* by dividing them by their standard error. We refer to these standard coefficients as 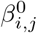 and the matrix of such standard coefficients for partitioning variables as the “standardized coefficient matrix”, 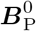.

A useful objective function for approximating the coefficient matrix with the contrast basis is the Euclidean norm of the projection of the standardized coefficient matrix onto contrast basis elements,

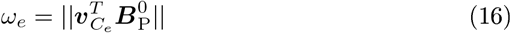

which captures the extent to which coefficients in 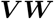 differ between the sets of species partitioned by the edge *e*. Coefficient contrasts are fast and easy to compute, but the algorithm described here minimizes the distance between ***VW*** and 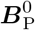. Other algorithms described below can more robustly identify the edge, *e*, whose reduced-rank approximation maximizes the likelihood.

#### Stepwise phylo factor contrasts

Other options for aggregation and contrast exploit the factor-contrasts built into generalized linear and additive modeling machinery. Factor contrasts using a variable `phylo` ∈ {*R, S*}, indicating which group a species is in, can capture the assumption of shared coefficients within-groups and contrast the coefficients between-groups in multivariate generalized linear modeling across all species. Stepwise, maximum-likelihood selection of`phylo` factor contrasts are a more accurate, yet computationally intensive, algorithm for partitioning exponential family random variables.

For example, a data frame contrasting how the counts of “birds” from “non-birds” react to meta-data *z*_2_ while controlling for *z*_1_ can be constructed as follows

| Site | Species | Abundance | $z_1$ | $z_2$ | phylo |
| --- | --- | --- | --- | --- | --- |
| 1 | Pigeon | 10 | 1 | .5 | $R$ |
| 1 | Dove | 8 | 1 | .5 | $R$ |
| 1 | Lizard | 1 | 1 | .5 | $S$ |
| 1 | Mouse | 3 | 1 | .5 | $S$ |
| 1 | Cat | 1 | 1 | .5 | $S$ |
| 2 | Pigeon | 2 | 0 | -2 | $R$ |
| 2 | Dove | 1 | 0 | -2 | $R$ |
| 2 | Lizard | 10 | 0 | -2 | $S$ |
| 2 | Mouse | 4 | 0 | -2 | $S$ |
| 2 | Cat | 3 | 0 | -2 | $S$ |
| ... | ... | ... | ... | ... | ... |

Phylofactorization can be implemented through a generalized linear model for a count family (e.g. Poisson, binomial, or negative binomial) using the formula

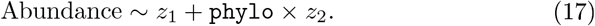

The `phylo` factor contrasts birds from non-birds and using its deviance as the objective function will find the edge *e*\* whose `phylo` factor maximizes the likelihood of the data.

In stepwise `phylo` factor contrasts, aggregation occurs within the likelihood function. The likelihood 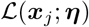 for a vector of binomial random variables ***x**_j_* can be written in exponential family form

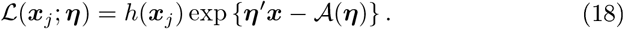

A two-factor model, such as ***x*** ~ `phylo`, will reduce the likelihood function from *s* parameters in ***η*** to two parameters, ***η*** ∈ (*η_R_, η_S_*), yielding

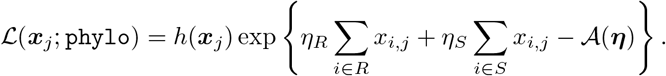

Aggregation, within the likelihood function above, is summation of data within-groups. Obtaining the maximum likelihood estimates, 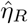 and 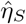, a contrast function can be defined as a difference of *η_R_* and *η_S_*, or test-statistic from the null hypothesis that *η_R_* = *η_S_*. For general purposes, the deviance of the `phylo` term in generalized linear or additive models serves as a useful contrast allowing one to identify the edge *e*\* whose `phylo`_*e*_ factor that maximizes the likelihood for the regression model containing the `phylo` factor.

Stepwise selection of maximum-likelihood `phylo` factor contrasts is a very accurate method for regression-phylofactorization of exponential family random variables. However, unlisting an entire dataset, computing a glm, and re-computing the glm for each edge in the phylogeny is computationally intensive.

#### Marginally Stable (mStable) Aggregation

Another option, aimed to allow maximum-likelihood estimation of `phylo` factor contrasts while reducing the computational diffculty, is to aggregate the data ***X*** prior to maximizing the likelihood in the generalized linear model. The method we present is to assume within-group homogeneity and aggregate exponential family random variables to a “marginally stable” exponential family random variable that can be used for downstream analysis. Marginal stability, to the best of our knowledge, has not been explicitly defined elsewhere, and thus we introduce the term here by loosening the definition of stable distributions [41].

#### Stable distribution

A distribution with parameters *θ*, 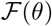, is said to be stable if a linear combination of two independent random variables from 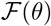 is also in 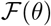, up to location and scale parameters.

#### Marginally stable distribution

A distribution with parameters {*θ*_1_, *θ*_2_}, 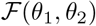, is said to be marginally stable on *θ*_1_ if 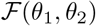 is it is stable conditioned on *θ*_1_ being fixed.

For example, the Gaussian distribution is stable: the sum of two Gaussian random variables is also Gaussian. Meanwhile, binomial random variables *Binom*(*ρ, N*) me marginally stable on *ρ*; random variables *x_i_* ~ *Binom*(*ρ, N_i_*) can be summed to yield *A*(***x***) ~ *Binom*(*ρ*, Σ *N_i_*). The marginal stability can also be used with transformations that connect the assumed distribution of the data to a marginally stable distribution. Log-normal random variables can be converted to Gaussians through exponentiation; chi random variables can be converted to chi-squared through squaring - random variables from many distributions may be analyzed by transformation to a stable or marginally stable family of distributions. Such transformation-based analyses implicitly define aggregation through a generalized *f*-mean

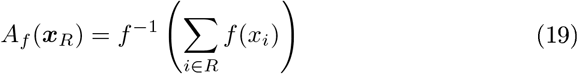

where *f*(*x*) = log(*x*) for log-normal random variables, *f*(*x*) = *x*^2^ for Chi random variables, etc. The goal of such aggregation, whether through exploiting marginal stability or generalized *f*-means or other group operations in the exponential family, is to produce summary statistics for each group, *R* and *S*, in a manner that permits generalized linear modeling of the summary statistics. By ensuring summary statistics are also exponential-family random variables, one can perform a factor-contrast style analysis as described above but only on the two summary statistics and not on all *s* species. Doing so can greatly reduce the computational load of phylofactorizing large datasets and, as we show below, can increase the power of edge-identifìcation even when the within-group homogeneity assumption does not hold. Marginal stability, for the purposes of phylofactorization, must be on the parameter of interest in generalized linear modeling (Figure 3a).

Marginal stability opens up more distributions to stable aggregation. Presence absence data, for instance, can be assumed to be Bernoulli random variables. The assumption of within-group homogeneity for the probability of presence, *ρ*, allows addition of Bernoulli random variables within each group, *R* and *S*, to yield a respective binomial random variable, *x_R_* and *x_S_*. Likewise, the addition of a set of binomial random variables with the same probability of success, *ρ*, yields an aggregate binomial random variable. A set of exponential random variables with the same rate parameter, *λ*, can be added to form a gamma random variable. Gamma random variables, *x_i_* ~ *Gamma*(*κ_i_, θ*), parameterized by their shape, *κ_i_*, and scale, *θ*, are marginally stable on *θ*. Addition of geometric random variables with the same rate parameter forms a negative binomial, and the addition of a set of negative binomial random variables, *x_i_* ~ *NB*(*π_i_, ρ*), with the same probability of success *ρ* but different numbers of failures, *π_i_*, can be aggregated into 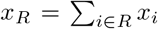 where 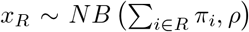. All of these distributions are not stable, but they are marginally stable.

Marginally stable aggregation can be made efficient by matrix multiplication onto one-vectors **1**_*R*_ and **1**_*S*_ whose ith entries are 1 for all *i* ∈ *R, S*, respectively, and 0 otherwise. Assuming a Poisson or negative binomial count model for the bird/non-bird data frame above, the data frame is reduced to and the same equation (17) can be used for phylofactorization through `phylo` factor-contrasts.

| Site | Species | Abundance | $z_1$ | $z_2$ | phylo |
| --- | --- | --- | --- | --- | --- |
| 1 | Bird | 18 | 1 | .5 | $R$ |
| 1 | Non-Bird | 5 | 1 | .5 | $S$ |
| 2 | Bird | 3 | 0 | -2 | $R$ |
| 2 | Non-Bird | 17 | 0 | -2 | $S$ |
| ... | ... | ... | ... | ... | ... |

Aggregation to a marginally stable distribution is computationally efficient but will only outperform maximum-likelihood estimation if the within-group heterogeneity is small. For 700 replicates for each combination of sample size *n* ∈ {5, 10, 30, 60}, effect size *δ* ∈ {0, 0.375, 0.75, 1.125, 1.5, 1.875, 2.25, 2.625, 3}, and within-group variance *σ* ∈ {0, 1, 2}, we simulated three explanatory variables {*z*_1_, *z*_2_, *z*_3_} as independent, identically distributed *n*-vectors of standard normal random variables. The log-odds of presence for individual *i* in group *R* or group *S* was modeled as

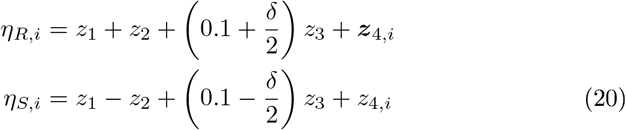

where 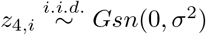 are independent Gaussian random variables particular to the individual and sample. The data were either kept as Bernoulli random variables or aggregated via summation to binomial random variables and then analyzed using factor contrasts in a generalized linear model of the form

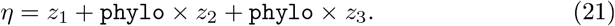

The objective function was the deviance from the final term, `phylo` × *z*_3_. The probability of identifying the correct edge and the distance between the identified and correct edge (in the number of nodes separating the two edges) are plotted in Figure 4b. The method of factor-contrasts for glm-phylofactorization asymptotically approaches perfect edge-identification, both in the probability of detecting the correct edge and in distance from the correct edge, as the sample sizes and effect sizes increase. Aggregation to binomial and subsequent factor-contrast of the aggregates slightly improved the power of edge-identification in these simulations. The improved accuracy of marginally-stable aggregation decreases with differences in within-group means, as opposed to an addition of individual within-group variance through ***z***_4,*i*_, as illustrated below. However, marginally-stable aggregation performs reasonably well and, crucially, scales well with increasing numbers of species and sample size. Consequently, if the datasets are large and the within-group homogeneity across samples is small, marginally-stable aggregation and stepwise construction of factor contrasts may be a useful tool for regression-phylofactorization of exponential family random variables.

**Figure 4:**
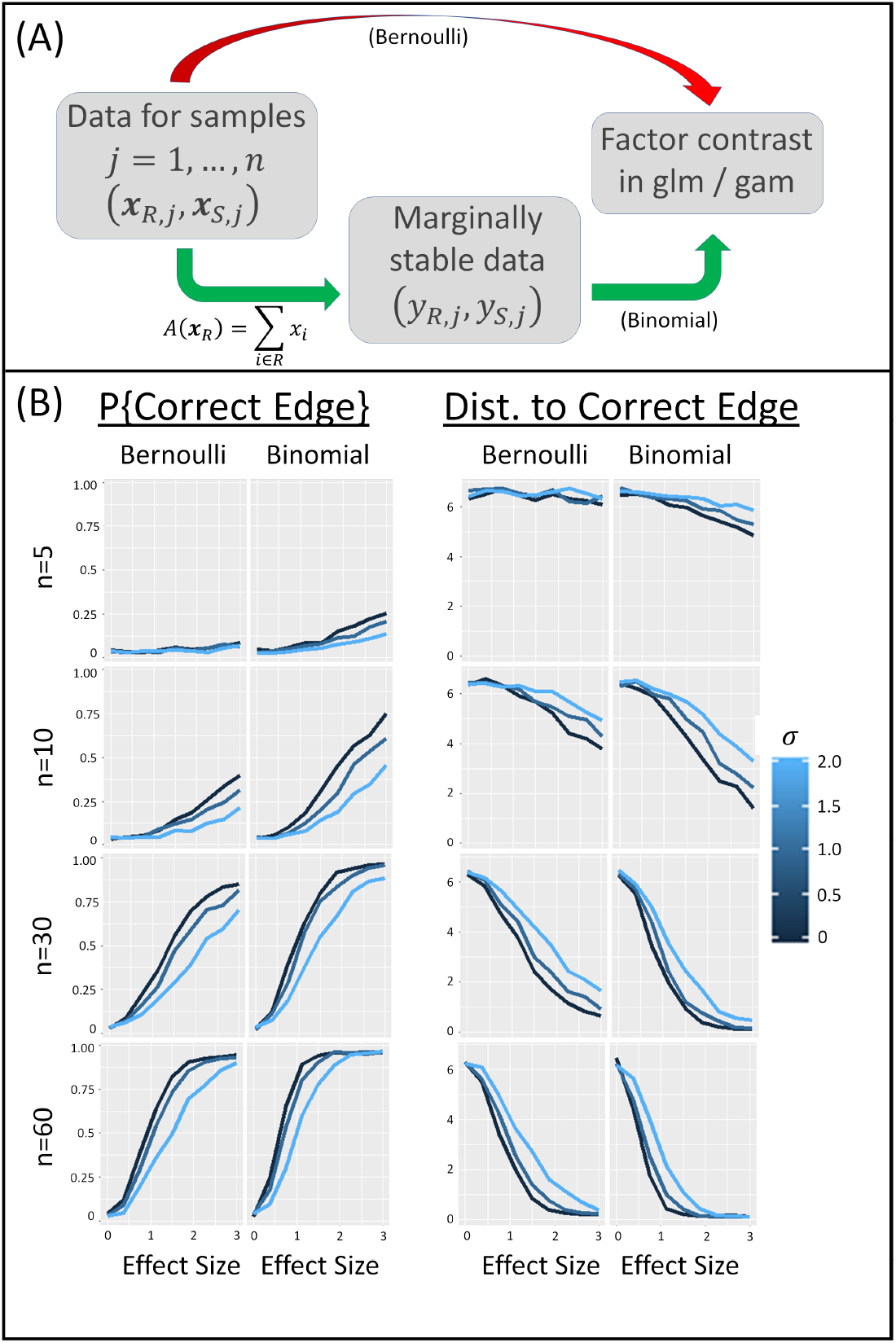
`phylo` factor contrasts can allow phylofactorization of exponential family random variables. (A) Each edge separates the species in a sample into two groups. These groups can be used as factors directly in a generalized linear model as in equations 17 and 21. Alternatively, a within-group homogeneity assumption can be used to aggregate data of many exponential family random variables to a marginally stable distribution, such as addition of Bernoulli random variables with the same probability of success to a binomial random variable. Regression on marginally stable random variables may dramatically reduce computational costs and, if within-group heterogeneity is low, improve accuracy. (B) Simulations of Bernoulli presence/absence data of 30 species with a random phylogeny suggest that aggregation to binomial improves power across a range of effect sizes, *δ*, (x-axis), sample sizes, *n* (rows), and within-group heterogeneity, *σ*. Here, aggregation of presence-absence data to binomial random variables for subsequent factor-contrasts outperformed the raw factor contrast of Bernoulli presence/absence data, suggesting it is at least a viable tool for large datasets. The generality of improved power of regression on surrogate, marginally stable aggregates remains to be seen.

#### Mixed Algorithm

Coefficient contrasts are computationally easy yet inac-curate, whereas stepwise `phylo` factor selection (without marginally-stable aggregation) is accurate yet computationally demanding (Figure 5). It’s possible to develop mixed algorithms with accuracy similar to stepwise `phylo` factor selection and reduced computational costs more similar to coefficient contrasts or marginally-stable aggregation. We present one example. In the first stages of the algorithm, multivariate generalized linear modelling is performed as for coefficient contrasts. For each iteration, coefficient contrasts (equation 16) are used to narrow down the set of possible edges, {*e*}_*top*_, to a set of edges with high objective functions from standardized coefficient contrasts. We use the top 20% of edges based on *ω_e_* in equation 16, resulting in an approximately 80% speed-up compared to the brute-force `phylo` factor contrast algorithm. For only these edges, `phylo` factors are considered and the winning edge is the top-quantile edge which maximizes the deviance of its `phylo` factor contrast.

**Figure 5:**
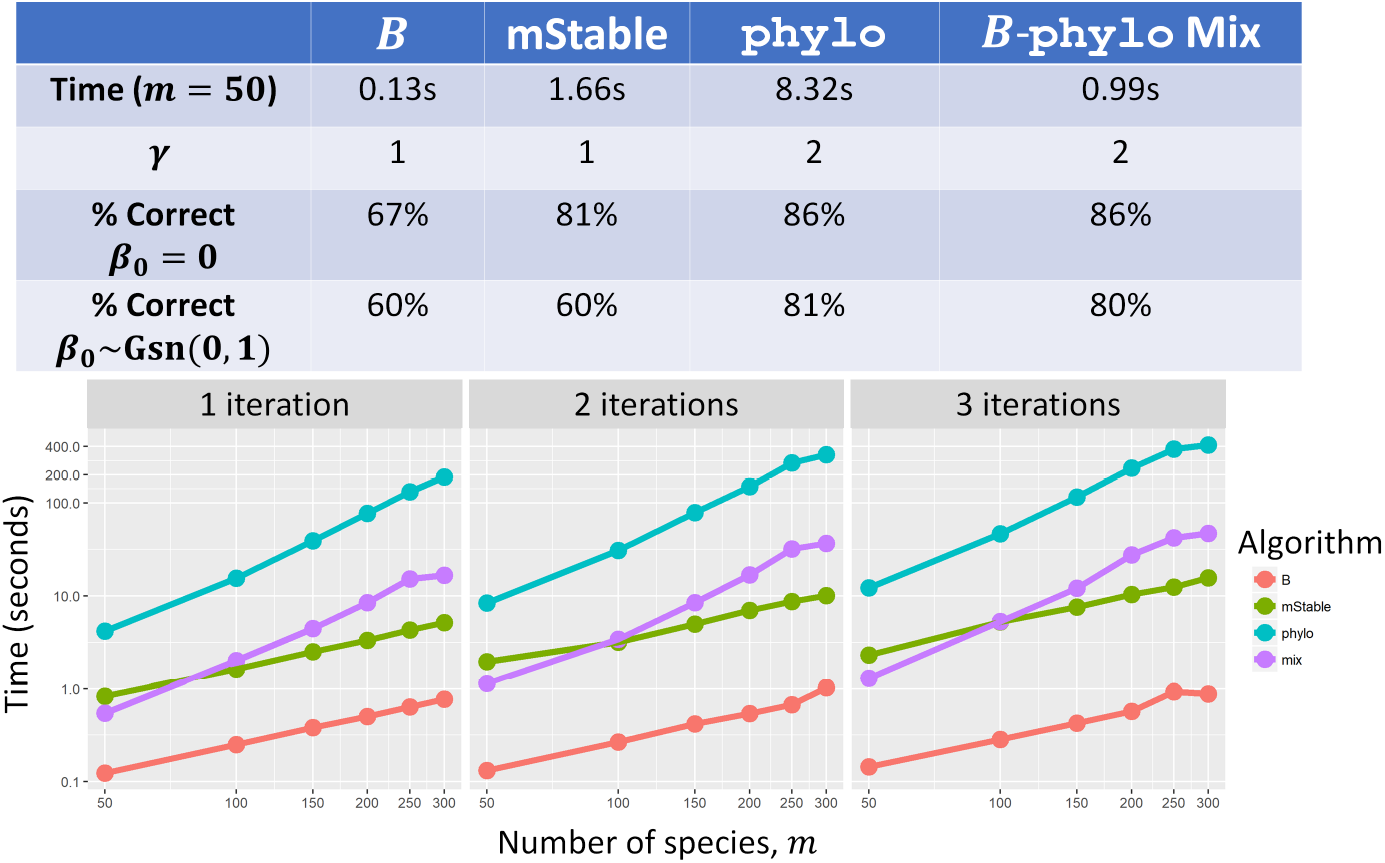
The accuracy, computation time and scaling of four algorithms for generalized phylofactorization. Algorithms are compared via the baseline time for two factors with *m* = 50 species, the scaling coefficient *γ* in *time* ∝ *m^γ^*, and percent of correctly identified edges in simulated data with *m* = 50 species and 2 affected clades. Stepwise `phylo` factor contrasts have high accuracy but are computationally costly and scale quadratically with the number of species. Marginally stable (mStable) aggregation scales linearly with *m* but only performs well when *β*_0_ = 0. Computation time can be reduced and accuracy preserved if coefficient contrasts in equation 16 are used to narrow the set of edges considered for rigorous phylo factor contrasts.

#### Algorithm comparison

We compare the performance of the four algorithms listed above by testing how well they can correctly identify the affected edges, 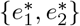, and how long they take to extract a variable number of factors. The four algorithms tested are:

- “B”: standardized coefficient contrasts
- “phylo”: Unaggregated `phylo` factor contrasts
- “mStable”: marginally-stable aggregation followed by `phylo` factor contrasts
- “mix”: Use of the “phylo” algorithm on only the top 20% of edges.

For edge identification, presence/absence data, *x_i,j_*, were simulated for a set of *s* = 50 species and *n* = 40 samples. The logit probability of all species was modelled as

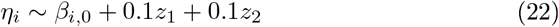

where 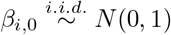 broke the within-group homogeneity in mean-probability of presence/absence. For comparison, the case with *β*_0,*i*_ = 0 for all species *i* is also considered. The other two explanatory variables, *z*_1_ and *z*_2_, were the partitioning variables differentiating species separated by edges. Two non-nested clades, one containing 21 species and the other containing 5 species, had a different association with the meta-data:

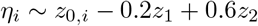

for *i* in either of the two affected clades. To add an additional level of complexity, the two meta-data variables were given multicolinearity by simulating *z*_1_ ~ *Gsn*(0, 1) and *z*_2_ ~ *Gsn*(*z*_1_, 1). The algorithms were run for two factors and the number of correctly identified edges (out of 2) was tallied across 1000 replicates (e.g. an algorithm that was 80% correct identified 1600 correct edges over 1000 replicates). The times for each of these algorithms to compute two factors was also recorded. To compare the scaling of the algorithms, null data were simulated across a range of species richness *m* ∈ {50, 100, 150, 200, 250, 300} and across a range of factors *t* ∈ {1, 2, 3}.

The stepwise `phylo` factor contrasts by maximizing the total deviance of `phylo` ∗ (*z*_1_ + *z*_2_) had the greatest accuracy but also the slowest computation time (Figure 5). The time required to compute `phylo` factor contrasts scale quadratically with the number species, *m*, whereas coefficient contrasts and marginally stable (mStable) aggregation scale linearly. Marginally stable aggregation only performs well when *β*_*i*,0_ = 0 for all species, *i*, and when the within-group heterogeneity is small. The accuracy of `phylo` factor contrasts can be preserved and the computation time reduced by selecting the top 20% of edges based on coefficient contrasts. The computation of multiple generalized linear models across edges can be parallelized to reduce computation time, and such parallelization is built into the R package `phylofactor`.

#### Summary of generalized phylofactorization

We have presented algorithms to perform regression-phylofactorization for non-Gaussian data. These algorithms can be called within the function `gpf`(). The stepwise selection of `phylo` factor contrasts is best able to correctly identify edges and is easily parallelizable. The computation time of stepwise `phylo` factor contrasts can be reduced by narrowing the set of considered edges to those with high coefficient contrasts. Marginally stable aggregation may be a promising alternative for faster algorithms as it scales linearly with the number of species, but marginally stable aggregation only performs well when there is little systematic difference in the mean, *β*_*i*,0_, across species, *i*.

There are fruitful avenues for future research to refine the algorithms for phylofactorization of big-data consisting of non-Gaussian exponential family random variables. These algorithms are intimately related to reduced rank regression and generalized linear modelling with shared coefficients. Reduced-rank regression considers a compact set of possible basis vectors and, consequently, can use gradient-descent methods to find maximum-likelihood estimates. The constrained set of allowable contrasts in the phylogeny precludes gradient-descent and produces problems directly analogous to those in phylogenetic components analysis and thus we have focused on explicit testing of all possible allowable contrasts in the phylogeny or, in the case of the mixed algorithm, testing a subset of contrasts believed to contain the winning edge, *e*\*. These methods can extend to generalized additive models and, as we discuss below, spatial and time-series data as well.

### Phylogenetic factors of space and time

Phylofactorization can also be used in analyses of spatial and temporal patterns. We’ve demonstrated phylofactorization through examples of cross-sectional data through two-sample tests, analyses of contrast-basis projections, and use of `phylo` factor contrasts in communities sampled across a range of meta-data. These same tools can be used for phylofactorization-based analysis of spatial and temporal ecological data. Samples of a community over space and time can be projected onto contrast basis elements and the resulting component scores, *y_e_*, can be analyzed much like PhyCA to identify the phylogenetic partitions of community composition over space and time. Spatial samples can be analyzed using `phylo` factor contrasts as defined for generalized linear models. Multivariate Autoregressive Integrated Moving Average (ARIMA) models can be constructed either as ARIMA models of the component scores, *y_e_*, or as multivariate ARIMA models with `phylo` factor contrasts as used in generalized linear models perform phylogenetic partitions based on differences in drift, volatility, and other features of interest. Coefficient matrices, including spatial and temporal autocorrelation matrices or coefficients for extrinsic meta-data ***Z***, can be approximated with phylogenetic contrast-bases as in equation (15).

Marginally stable aggregation in spatial and temporal data requires a more complex consideration of the marginal stability of spatially explicit random variable and stochastic processes. “Stability”, for spatially and temporally explicit random variables, must preserve the underlying model for the spatial or temporal process being used for analysis. An example of a less obvious marginally stable aggregation of time-series data is the stability of neutral drift (sensu Hubbell [22]) to grouping.

Neutral communities fluctuate, and those fluctuations have a drift and volatility unique to neutral drift. Neutral drift can also be defined either by discrete, finite-community size urn processes or stochastic differential equations for the continuous approximations of finite but large communities. Recently, Washburne et al. [50] articulated the importance of a feature of neutral drift which enables time-series neutrality tests: its invariance to grouping of species. If a stochastic process of relative abundances, ***X**_t_*, obeys the probability law defined by neutral drift, then any complete, disjoint groupings of ***X**_t_* also obeys the probability law for a lower-dimensional neutral drift. Thus, neutral processes are stable to aggregation by grouping or summation of relative abundances. Collapsing all species into two disjoint groups, *R* and *S*, yields a two-dimensional neutral drift with a well-defined neutrality test for time-series data. Specifically, if ***X**_t_* is a Wright Fisher process and *R* and *S* are disjoint groups whose union is the entire community, the quantity

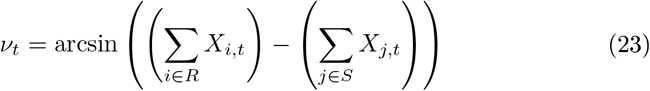

has a constant volatility which can be used to define a neutrality test for time-series data. Thus, phylofactorization can be done to partition edges across which the dynamics appear to be the least neutral. For the test developed by Washburne et al., the aggregation operation is the *L*_1_ norm and the contrast operation is subtraction:

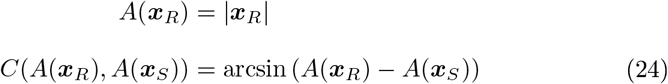

and the objective function, *ω*, for edge *e* is the test-statistic of a homoskedasticity test of *C_e_*. Neutrality is a relative measure - biological units are neutral relative to one-another - and thus the use of aggregation of species into a unit and a contrast of two units is a natural connection between the theory and operations of phylofactorization and the concept of neutrality.

### Statistical Challenges

We present a unifying algorithm which partition organisms into functional groups by identifying meaningful differences or contrasts along edges in the phylogeny. Phylofactorization is formally defined as a graph-partitioning algorithm. However, maximizing the variance of the data projected onto contrast basis elements corresponding to edges in the phylogeny is a constrained principal components analysis. The use of regression-based objective functions and the iterative construction of a low-rank approximation of a data matrix is similar to factor analysis. The discovery of a sequence of orthogonal factor contrasts in generalized linear models is a form of stepwise/hierarchical regression and partitioning a coefficient matrix ***B*** is a reduced-rank regression method. The maximization of the objective function at each iteration is a greedy algorithm. Each of these connections between phylofactorization and other classes of methods produces a body of literature from related methods which could inform phylofactorization and facilitate rapid development of this exploratory tool into a more robust, inferential one.

There are statistical challenges common across many methods for phylofactorization. In this section, we enumerate some of the statistical challenges and discuss work that has been done so far. First, as with any method using the phylogeny as a scaffold for creating variables or making inferences, the uncertainty of the phylogeny and the common use of multiple equally likely phylogenies warrant consideration and further method development. Other challenges discussed here are: understanding the propagation of error; development of Metropolis algorithms to better arrive at global maxima; the appropriateness, and error rates, of phylofactorization under various evolutionary models underlying the effects (e.g. trait differences, habitat associations, etc.) and residuals in our data; understanding graph-topological biases and confidence regions; cross-validating the partitions and inferences from phylofactorization; determining the appropriate number of factors and stopping criteria to stop a running phylofactorization algorithm; and understanding the null distribution of test-statistics when objective functions being maximized are themselves test-statistics from a well-known distribution. Any exploratory data analysis tool can be made into an inferential tool with appropriate understanding of its behavior under a null hypothesis, and the connections of phylofactorization to related methods can accelerate the development of well-calibrated statistical tests for phylogenetic factors.

#### Phylogenetic inference

So far we have assumed that the phylogeny is known and error free, but the true evolutionary history is not known - it is estimated. Consequently, phylofactorizations are making inferences on an uncertain scaffold; the more certain the scaffold, the more certain our inferences about a clade. Two challenges remain for dealing with phylofactorization on an uncertain phylogeny. For a consensus tree, there is the question of what statistics of the consensus are most easily integrated for precise statements of uncertainty in phylofactorization inferences. Bootstrapped confidence limits for monophyly [12] are the most commonly used statement of uncertainty for a consensus tree, but there may be others as well. Different organisms will have different leverages in regression or two-sample test phylofactorization, and thus monophyly is only part of the picture: leverage is another. For a set of equally likely bootstrapped trees, there is a need to integrate phylofactorization across trees. Phylofactorization of sets of equally likely phylogenies has not yet been done, but may be a fruitful avenue for future research. One last option for researchers with trees containing clades with low bootstrap monophyly is to lower the resolution of the tree. Phylofactorization can still be performed on a tree with polytomies - the mammalian phylogeny used above contained many - and reducing the number of edges considered at each iteration can focus statistical effort (and chances of false-discovery) on clades about which the researcher is more certain.

#### Propagation of error

Phylofactorization is a greedy algorithm. Like any greedy algorithm, the deterministic application of phylofactorization is non-recoverable. Choosing the incorrect edge at one iteration can cause error to propagate, potentially leading to decreased reliability of downstream edges. Little research has been done towards managing the propagation of error in phylofactorization, but recognizing the method as a greedy algorithm suggests options for improving performance. Stochastic-optimization schemes, such as replicate phylofactorizations using Metropolis algorithms and stochastic sampling as implemented in the mammalian tree phylofactorization (sampling of edges with probabilities increasing monotonically with *ω_e_* and picking the phylofactor object which maximizes a global objective function), may reduce the risk of error cascades in phylofactorization [20].

#### Behavior under various evolutionary models

Phylofactorization is hypothesized to work well under a punctuated-equilibrium model of evolution or jump-diffusion processes [15, 26] in which jumps are infrequent and large, such as the evolution of vertebrates to land or water. If few edges have large changes in functional ecological traits underlying the pattern of interest, phylofactorization is hypothesized to work well. Phylofactorization may also work well when infrequent life-history traits arise or evolutionary events occur (such as ecological release) along edges and don’t yield an obvious trait but instead yield a correlated, directional evolution among descendants. Phylofactorization of mammalian body sizes yielded a scenario hypothesized to be in this category. In this case the exact trait may not have arisen along the edge identified, but a precursor trait, or a chance event such as extinctions or the emergence of novel niches, may precipitate downstream evolution of the traits underlying phylofactorization. Both aggregation and contrast functions can incorporate phylogenetic structure and edge lengths to partition the tree based on likelihoods of such evolutionary models. The sensitivity of phylofactorization to alternative models, such as continuous Brownian motion and Ornstein-Uhlenbeck models commonly used in phylogenetic comparative methods [13, 19], remains to be tested and will likely vary depending on the particular method used.

#### Basal/distal biases

Researchers may be interested in the distribution of factored edges in the tree. If a dataset of microbial abundances in response to antibiotics is analyzed by regression-phylofactorization and results in many tips being selected, a researcher may be interested in quantifying the probability of drawing a certain number of tips given t iterations of phylofactorization. Alternatively, if several edges are drawn in close proximity researchers may wonder the probability of drawing such clustered edges under a null model of phylofactorization. For another example, researchers may wonder if the number of important functional ecological traits arose in a particular historical time window (e.g. due to some hypothesis of important evolutionary event or environmental change), and thus want to test the probability of drawing as many or more edges than observed under a null model of phylofactorization. All of these tests would require an accurate understanding of the probability of drawing edges in different locations of the tree.

All methods described here, save the Fisher exact test, have a bias for tips in the phylogeny (Figure 6). Such biases affect the calibration of statistical tests of the location of phylogenetic factors, such as a test of whether/not there is an unusually large number of differentiating edges in mammalian body mass during or after the K-Pg extinction event.

**Figure 6:**
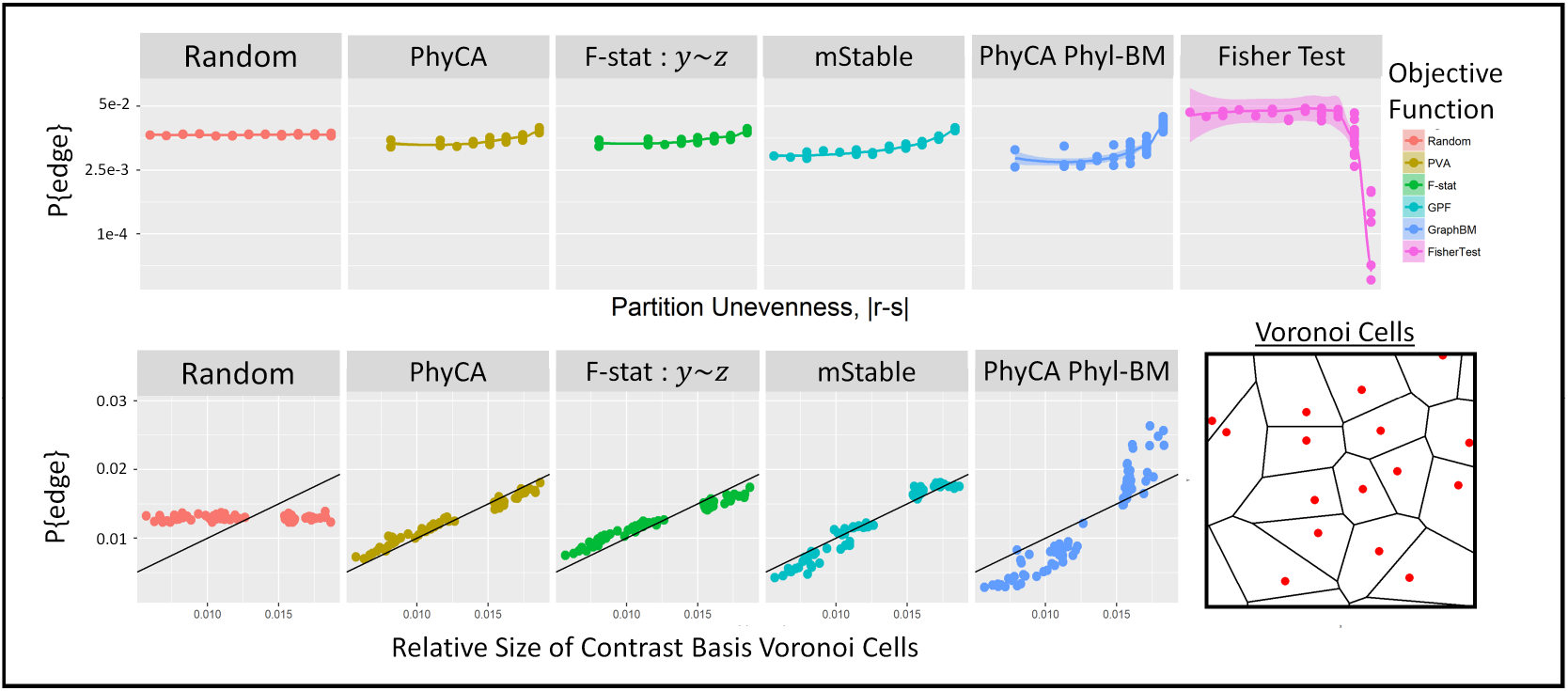
Graph topological bias in null data and the relative size of Voronoi cells of contrast basis elements. The method and the null distribution of the data determine graph-topological bias of phylofactorization. A random draw of edges does not discriminate against edges based on the relative sizes of two groups contrasted by the edge, but 16,000 replicate phylofactorizations of null data reveal that contrast-basis methods are slightly biased towards uneven splits (e.g. tips of the phylogeny). Standard Gaussian null data were used for PhyCA, F-statistics from regression on contrast basis elements (*y_e_* ~ *z*), and binomial null data was used for generalized phylofactorization (gpf) through marginally-stable aggregation. Other methods, such as Fisher’s exact test of a vector of Bernoulli random variables, have opposite biases. The tip-bias of contrast-basis analysis is amplified for marginal-stable aggregation in generalized phylofactorization, and amplified even more if the null data have residual structure from a Brownian motion diffusion along the phylogeny (Phyl-BM). The common bias when using contrast bases across a range of objective functions is related to the uneven relative sizes of Voronoi cells produced by the bases, simulated here by equation (25).

Phylofactorization using the contrast basis is biased towards the tips of the tree. Some progress can be made towards understanding the source of basal/distal biases in phylofactorization via the contrast-basis. The biases from analyses of contrast basis coordinates, ***y**_e_*, stem from a common feature of the set of *K_t_* candidate basis elements 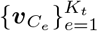 considered at iteration *t* of phylofactorization. For the example of the t-test phylofactorization of a vector of data, ***x*** the winning edge *e*\* is

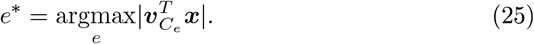

If all basis elements have unit norm, which they do under equation (5), then each basis element being considered corresponds to a point on an *m*-dimensional unit hypersphere. If the data, ***x***, are drawn at random, such that no direction is favored over another, the probability that a particular edge *e* is the winning edge is proportional to the relative size of its Voronoi cell on the surface of the unit *m*-hypersphere. Thus, the basal/distal biases for contrast-basis analyses with null data assumed to be drawn from a random direction can be boiled down to calculating or computing the relative sizes of Voronoi cells. For our simulation, we estimated the size of Voronoi cells through matrix multiplication

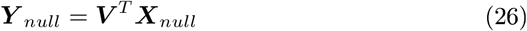

were ***V*** is a matrix whose columns *j* is the contrast basis elements for edge *e_j_* being considered and ***X**_null_* is the dataset simulated under the null model of choice whose columns are independent samples ***x**_j_*. Each column of ***Y**_null_* contains the projections of a single random vector - the element of each column with the largest absolute value is the edge closest to that random vector.

#### Graph-topology and confidence regions

As a graph-partitioning algorithm, phylofactorization invites a novel description of confidence regions over the phylogeny. The graph-topology of our inferences - edges, and their proximity to other edges, both on the phylogeny and in the *m*-dimensional hypersphere discussed above - can be used to refine our statements of uncertainty. 95% Confidence intervals for an estimate, e.g. the sample mean, give bounds within which the true value is likely to fall 95% of the time in random draws of the estimate. Confidence regions are multi-dimensional extensions of confidence intervals. Conceptually, it’s possible to make similar statements regarding phylogenetic factors - confidence regions on a graph indicating the regions in which the true, differentiating edge is likely to be.

Extending the concept of confidence regions to the graph-topological inferences from phylofactorization requires useful notions of distance and “regions” in graphs. One example of such a distance between two edges is a walking distance: the number of nodes one crosses along the geodesic path between two edges. Alternatively, one could define regions in terms of years or branch-lengths. Defining confidence regions in phylofactorization must combine the uneven Voronoi cell sizes as well as the geometry of the contrast basis. For low effect sizes, confidence regions extend to distant edges on the graph whose contrast basis have a large relative Voronoi cell size (e.g. the tips). As the effect sizes increase, confidence regions over the graph are better described in terms of angular distances between the contrast basis elements and that of the winning edge, *e*\* (Figure 7).

**Figure 7:**
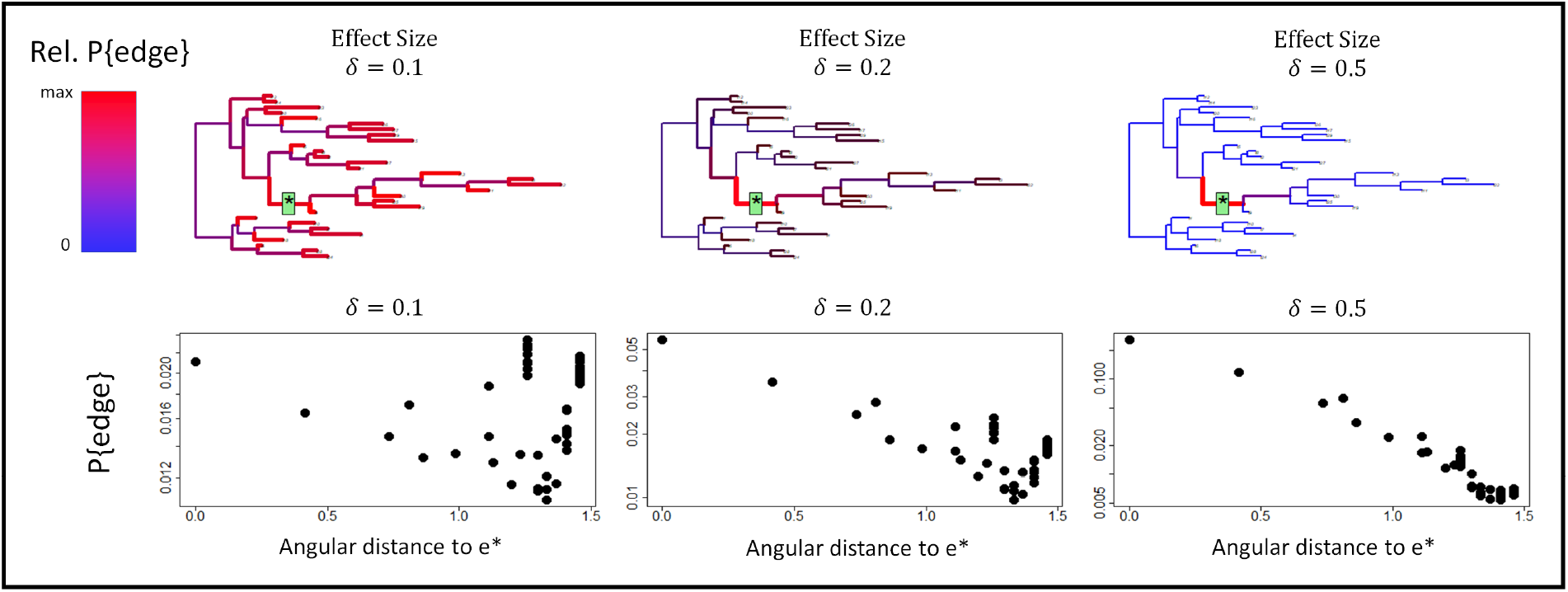
Graph-topological confidence regions for phylofactorization. **C**onfidence regions around inferred edges must use distances relevant to the method and graph topology. A tree with 30 species was given a fixed effect about edge *e*\* in their mean values as a function of meta-data *z* ~ *Gsn*(±*δ*/2, 1) 7 × 10^5^ iterations of phylofactorization were run and the relative probability of drawing each edge was visualized through both the color and width of the edge. The relationship between the angular distance of an edge’s contrast basis element to that of *e*\* and the probability of drawing the edge indicate that for low effects, confidence intervals must incorporate a mix of tip-bias and angular distance, but larger effect sizes, in which the edge drawn is reliably in the neighborhood of *e*\*, the angular distance of contrast basis elements capture confidence regions around the location of inferred phylogenetic factors.

#### Cross-validation

How do we compare phylofactorization across datasets to cross-validate our results? If a researcher observes a pattern in the ratio of squamates to mammalian abundances in North America, say a decrease in the ratio of lizard and snake to mammal abundance with increasing altitude, they may wish to cross-validate their findings in other regions, including regions with few or none of the same species in the original study. Researchers replicating the study in Australia and New Zealand would have to grapple with whether or not to include monotremes in their grouping of “mammals” and whether or not to include the tuatara, a close relative of squamates, in their grouping of “squamates” - such branches were basal to the squamate & mammalian clades contrasted in the hypothetical North American study.

Phylofactorization formalizes the issues arising with such phylogenetic cross-validation (Figure 8). If all species in the training/testing datasets can be located on a universal phylogeny, phylofactorization of a training set of species and data identifies edges or links of edges in the training phylogeny which are guaranteed to correspond to edges or links of edges in the universal phylogeny. The testing set of species may introduce new edges to the phylogeny which interrupt the links of edges in the universal phylogeny along which training contrasts were conducted. In the example above, the tuatara and monotremes all interrupt the link of edges separating North American mammals from North American reptiles on the universal phylogeny.

**Figure 8:**
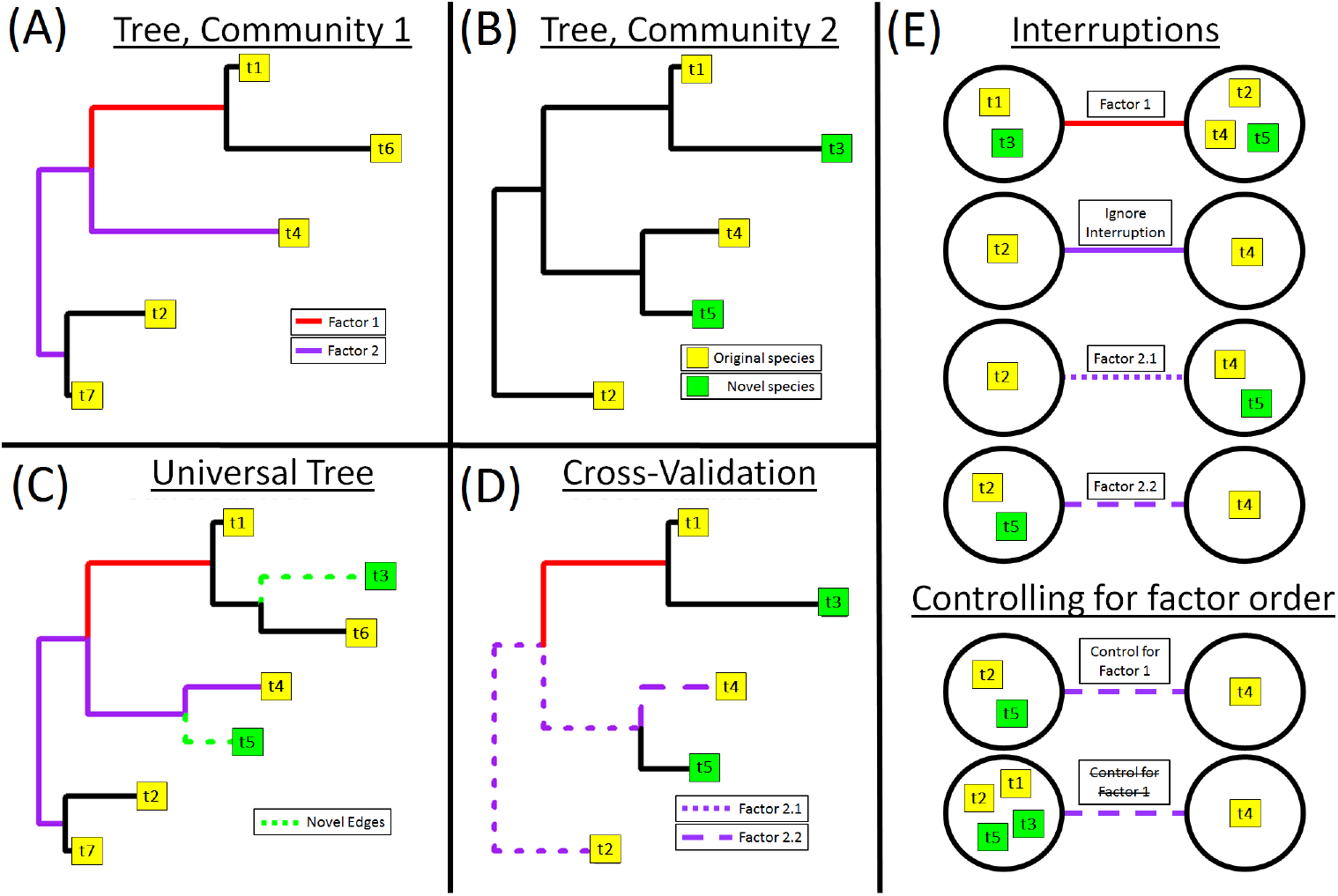
Graph-topological considerations with cross-validation. **(A)** The training community has 5 species (yellow boxes) split into two factors. The second factor forms a partition separating t4 from {t2,t7}. The second factor does not correspond to a single edge, but instead a chain of two edges. **(B)** A second, testing community is missing species t6 and t7 and contains novel species t3 and t5 (green boxes). **(C)** All factors can be mapped to chains of edges on a universal phylogeny. Novel species “interrupt” edges in the original tree; cross-validation requires deciding what to do with novel species and interrupted edges. Species t3 does not interrupt a factored edge, and so t3 can be reliably grouped with t1 in factor 1. However, species t5 interrupts one of the edges in the edge-path of factor 2. **(D-E)** Interruptions can be ignored, or they can be used to refine the location of important edges (illustrated in Factor 2.1 and Factor 2.2). Another topological and statistical question is whether/not to control for factor order. For instance, controlling for factor order with Factor 2.2 would partition t4 from {t2,t5}. Not controlling for factor order would partition t4 from {t1,t2,t3,t5}.

Robust cross-validation for phylofactorization requires directly addressing the issues arising from the interruptions of edges produced by novel species. Interruptions may be either ignored, or used to refine the inference. Returning to the previous example, one can use the presence of monotremes and tuatara to refine the definition of North American mammals to mean “all mammals” and “all placental and marsupial mammals”, and likewise one can optionally refine the definition of “squamates” to the broader “Lepidosauria” clade.

#### Stopping Criteria

Often, it’s desireable to obtain a minimal set of partitions to prioritize findings, simplify high-dimensional data, and focus effort on more certain inferences. Doing so requires a method for stopping phylofactorization. There are two broad options for stopping phylofactorization: a stopping function demonstrated to be sufficiently conservative, and null simulations allowing quantile-based cutoffs (e.g. stop phylofactorization when the percent variance explained by PhyCA is within the 95% quantile of null phylofactorizations). Null simulations may allow statistical statements stemming from a clear null model, but stopping criteria can be far more computationally efficient and can be constructed to be conservative.

Washburne et al. [51] proposed a stopping criterion for regression phylofactorization which extends to all methods of phylofactorization using an objective function that is a test-statistic whose null-distribution is known. The original stopping criterion is based on the fact that, if the null hypothesis is true, the distribution of P-values from multiple hypothesis tests is uniform. Phylofactorization performs multiple hypothesis tests at each iteration. At each iteration, one can perform a one-tailed KS test on the uniformity of the distribution of the P-values from the test-statistics on each edge; if the KS-test is non-significant, stop phylofactorization. KS-test stopping criteria can conservatively stop simulations at the appropriate number of factors when there is a discrete subset of edges with effects. Such a method performs similarly to Horn’s stopping criterion for factor analysis [21], whereby one stops factorization when the scree plot from the data crosses that expected from null data (figure 9). It’s also possible to first use a stopping criterion and subsequently run null simulations to understand the likelihood of observed results under a null model of the researcher’s choice (figure 9).

**Figure 9:**
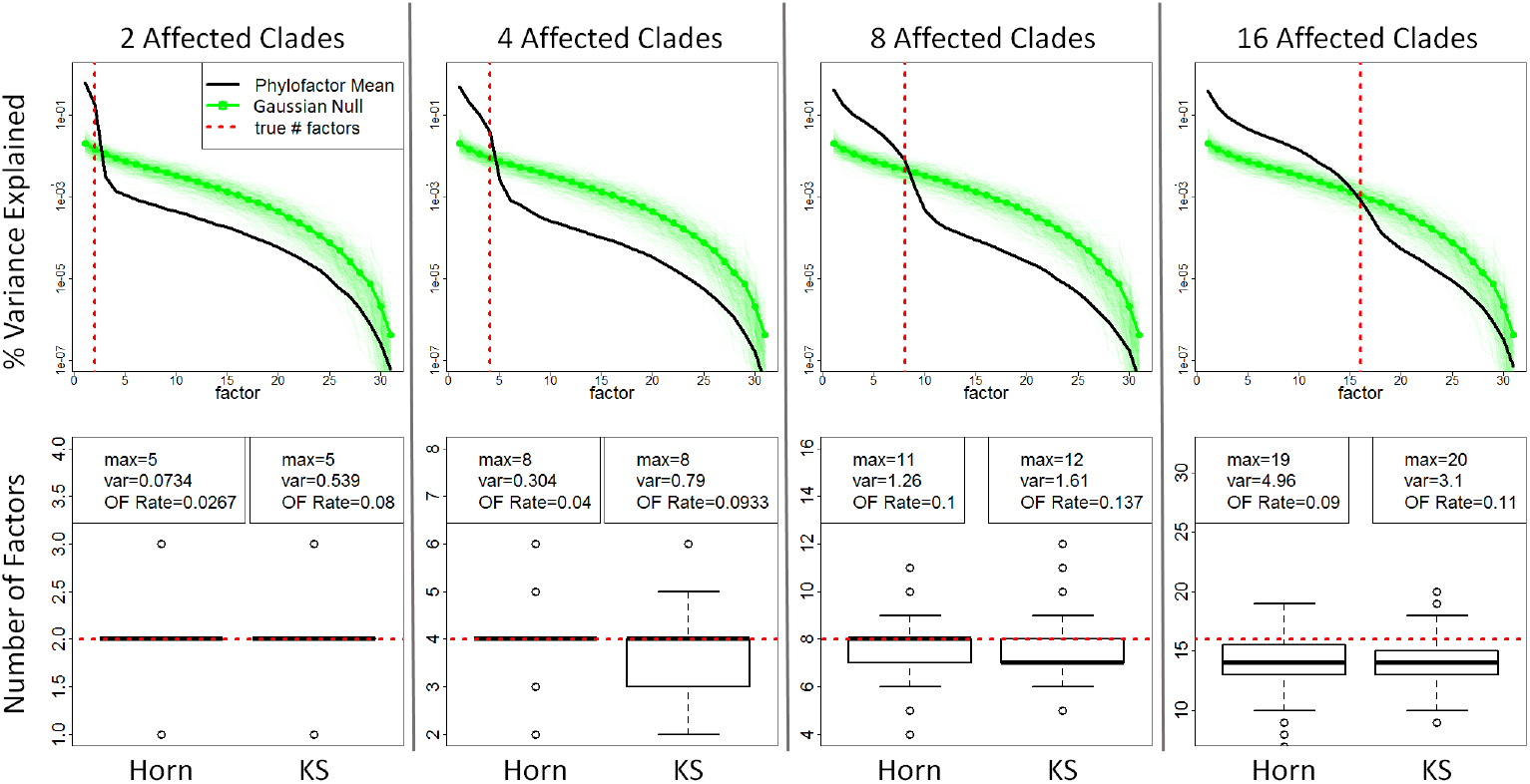
Null simulations and stopping criteria. A challenge of phylofactorization is determining the number of factors, *K*, to include in an analysis. Null simulations allow quantile-based cutoffs such as those in Horn’s parallel analysis from factor analysis. Stopping criteria stop phylofactorization using features available during phylofactorization of the observed data. Abundances of *m* = 32 species across *n* = 10 samples were simulated as i.i.d. standard Gaussian random variables. A set of *u* clades were associated with environmental meta-data, ***z***, where 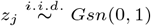. Regression-phylofactorization on the contrast-basis scores *y_e_* was performed on 300 datasets for each *u* ∈ {2, 4, 8, 16} and on data with and without effects. The objective function was the total variance explained by regression *y_e_* ~ *z*. (**top row**) The percent of the variance in the datseta explained at each factor (EV) decreases with factor, *t*, and the mean EV curve for data with *u* affected clades intersects the mean EV curve for null data near where *t* = *u*, motivating a stopping criterion (Horn) based on phylofactorization of null datasets, (**bottom row**) The Horn stopping criterion has a slightly lower over-factorization (OF) rate than the standard KS stopping criterion (where OF rate is the fraction of the 300 phylofactorizations of data with simulated effects in which *t* > *u*). However, the algorithms were not extremely different and both criteria can be modified to be made more conservative. The KS stopping criterion is far less computationally intensive for large datasets as it requires running phylofactorization only once. Null simulations, however, can allow inferential statistical statements regarding the null distribution of test statistics in phylofactorization.

#### Calibrating Statistical Tests for *ω_e*_*

Often, the objective function for the winning edge in phylofactorization, *ω_e*_*, corresponds directly to a common test-statistic. Applying a standard test for the resultant test-statistic, however, will lead to a high false-positive rate and an over-estimation of the significance of an effect, as the statistic was drawn as the best of many. Even when using a test-statistic not equal to the objective function, researchers should be cautious of dependence between their test-statistic and the objective function as a possible source of high false-positive rates. Two nmethods for calibrating, or making conservative, statistical tests of *ω_e*_* are multiple-comparisons corrections to control a family-wise error rate (or other multiple-hypothesis-test methods) or conservative bounds on the distribution of the maximum of many independent, identically distributed statistics. For example, if each edge of *K_t_* edges considered at iteration *t* sresulted in an independent *F*-statistic, *F_e_*, then the distribution of the maximum *F*-statistics, *F_e*_*, is

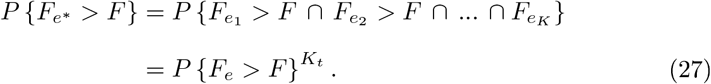

Such an approximation may be used to yield conservative estimates, but the *F*-statistics are not independent and thus more nuanced analyses are needed for well-calibrated statistical tests. More research is needed to obtain conservative bounds on test-statistics in phylofactorization.

#### Summary of limitations

Phylofactorization can be a reliable statistical tool with a careful understanding of the statistical challenges inherent in the method and shared with related methods such as graph-partitioning, greedy algorithms, factor analysis, and the use of a constrained, biased basis for matrix factorization. Phylofactorization can first and easiest be an exploratory tool, but all exploratory tools can be made inferential with suitable understanding of their behavior under an appropriate null model. For example, principal components analysis was and still is primarily an exploratory tool, but the discovery of the Marcenko-Pastur distribution [30] has improved the calibration of statistical tests on principal components for standardized, mean-centered data. Improved understanding of how uncertainties in phylogenetic inference translate to uncertainties in phylofactorization, conservative stopping criteria, null distributions of test-statistics for winning edges, propagation of error and stochastic sampling algorithms to avoid deterministic ruts, graph-topological biases and confidence regions on a graph, can all improve the reliability of phylofactorization as an inferential tool.

While phylofactorization was built with an evolutionary model of punctuated equilibria in mind, it may also work well under other evolutionary models such as correlated evolution among descendants of an edge. There are also many evolutionary models under which phylofactorization does not perform well. For instance the graph-topological biases of PhyCA are increased under a Brownian motion model of evolution. All statistical tools operate well under appropriate assumptions, and understanding the assumptions, as well as the known limitations, are necessary for responsible and academically fruitful use of statistical tools like phylofactorization.

## Discussion

Functional ecological traits underlie many observed patterns in ecology, including species abundances, presence/absence of species, and responses of traits or abundances to experimental conditions or along environmental gradients. Where the ecological pattern of interest is associated with heritable traits, the phylogeny provides a scaffold for the discovery of functional groupings of clades underlying the ecological pattern of interest. Traits arise along edges, and contrasting taxa on opposing sides of an edge allows one to uncover edges best separating species with different functional associations or links to the ecological pattern. By noting that each edge partitions the phylogeny into two disjoint sets of species, by generalizing the operations of “grouping” - aggregating and contrasting disjoint sets of species - and by defining the objective function of interest (the pattern), we have proposed a universal method for identifying relevant phylogenetic scales in ecological datasets.

Phylofactorization is a graph-partitioning algorithm intended to separate the phylogeny into binned phylogenetic units with a combination of high within-group similarity and high between-group differences. Two-sample tests are a natural method for making such partitions in vectors of data; such partitions can also be made with ancestral state reconstruction. The quantities used in two-sample tests can be extended to larger, real-valued datasets by analysis of a contrast basis. Objective functions for choosing the appropriate contrast basis include maximizing variance - a phylogenetic analog of principal components analysis - maximizing explained variance from regression, maximizing F-statistics from regression, and more. By partitioning coefficient matrices and using `phylo` factor contrasts, phylofactorization can be extended to generalized linear models, generalized additive models, and analyses of spatial and temporal patterns in ecological data.

We’ve illustrated that two-sample tests can partition a dataset of mammalian body mass into groups with very different average body masses. We’ve demonstrated that maximizing variance of data projected onto a contrast basis can identify major clades of bacteria in human feces that have been known, at a coarser resolution, to be highly variable and determined that one of the top phylogenetic factors in the American Gut dataset is a clade of Gammaproteobacteria associated with IBD and used recently in an effort to diagnose patients with Crohn’s disease. We’ve shown that analyses of contrast bases can use nonlinear regression, and within minutes of analysis on a laptop found a natural way put over 3,000 species into 5 binned phylogenetic units, sort them along an axis of the dominant explanatory variable, and produce a simplified story of how community composition changes in Central Park soils.

One can also perform phylofactorization when doing maximum-likelihood regression of exponential family random variables. The coefficient matrix can be approximated using the contrast basis, resulting in a phylogenetically-interpretable reduced-rank regression. Alternativley, it's possible to use `phylo` factor contrasts for a shared-coefficients model and maximum-likelihood based selection of edges for partitioning. One can either perform the factor contrasts on the raw data, or, for many exponential family random variables, one can aggregate the data from each group to a marginally stable distribution for more computationally efficient factor contrasts. These methods can be extended to spatial and temporal data. All methods discussed here can be implemented with the R package “phylofactor”, and scripts for running all analyses in this paper are available in the supplemental materials.

As with any method, there are limitations to be aware of. First, the general problem of separating species into *k* bins that maximize a global objective function is an NP hard problem. Second, like any greedy algorithm, purely deterministic phylofactorization may fall into ruts and errors in one step might propagate into downstream inferences. Third, the null distribution of test-statistics resulting from phylofactorization is not known; the resultant test statistics are biased towards extreme values. Null simulations, conservative stopping functions, and/or extremely stringent multiple comparisons corrections can be used to make inferences through phylofactorization while maintaining conservative bounds in family-wise error or false-discovery rate. When the objective function being maximized is also a test-statistic with a well-defined null distribution, one-sided KS-tests of the P-values from the test-statistic can serve as a computationally efficient and conservative stopping function. Fourth, common objective functions using the contrast basis will be biased due to the unequal relative sizes of the Voronoi cells of the contrast basis elements in the unit hypersphere in which they lie, with contrast basis elements corresponding to tips of the phylogeny tending to have larger relative Voronoi cell size than contrast basis elements corresponding to interior edges. Understanding the graph-topology of errors can assist the description of graph-topological confidence regions for each inference. Finally, phylofactorization formalizes the logic and challenges of cross-validating ecological comparisons even when the training and testing sets of species are completely disjoint. Many of these limitations may be resolved with future work, allowing the general algorithm and its common implementations to become a reliable, well-calibrated inferential tool.

Phylofactorization can objectively identify phylogenetic scales for ecological big-data and instantly produce avenues for future naturaly history research. By iteratively identifying clades, phylofactorization provides a sequence of low-rank approximations of a dataset, such as that visualized in figure 3c, which correspond to groups of species with a shared evolutionary history. What traits characterize the Chloracidobacteria which don’t like acidic soils? What traits characterize the monophyletic clade of Gammaproteobacteria that are associated with IBD? What traits underlie the Clostridia/Erysipelotrichi being such variable species in the American gut? The low-rank approximations of ecological data obtained by phylofactorization motivate subsequent questions best answered by life history comparisons, comparative genomics, microbial physiological studies, and other avenues of future research contrasting the species partitioned.

### Relation to other phylogenetic methods

Phylofactorization is proposed amidst an explosion of literature in phylogenetic comparative methods and various other phylogenetic methods for analyzing ecological datasets [29, 38, 14], and some careful thinking is beneficial to clarify the distinctions between the myriad methods.

Phylogenetic generalized least squares [16] aims to control for residual structure in the response variable expected under a model of trait evolution, and is thus used when performing regression on a trait, whereas phylofactorization aims to partition observed trait values or abundances into groups, separated byedges, with different means or associations with meta-data. Thus, while methods of phylogenetic signal, such as Pagel’s *λ* [35] or Blomberg’s *κ* [5], summarize global patterns of phylogenetic signal by parameterizing the extent to which a particular model of evolution can be assumed to underlie the residual structure of observed traits (often for downstream use in PGLS), phylofactorization iteratively identifies precise locations of putative changes and precise locations partitioning phylogenetic signal or structure.

Phylofactorization can be implemented by a contrast of ancestral state reconstructions of nodes separated by edges, for example by looking for edges with nodes whose reconstructed ancestral states are most different, but is limited by disallowing the descendant clade of an edge to impact the ancestral state of the edge’s basal node - a proper non-overlapping contrast would separate the groups of species being used to reconstruct each node, and thus phylofactorization can be implemented with ancestral state reconstruction under the assumption of time-reversible evolutionary models.

Phylogenetically independent contrasts [13] produces variables corresponding to contrasts of descendants from each node, whereas phylofactorization uses contrasts of species separated by an edge, picks out the best edge, splits the tree, and repeats. Phylofactorization develops a set of variables and an orthonormal basis to describe ecological data, but limits itself to bases interpretable as non-overlapping contrasts along edges; eigenvectors of phylogenetic distances matrices or covariance matrices under diffusion models of traits [35], are not encompassed in phylofactorization as they do not construct non-overlapping contrasts along edges. Such eigenvector methods construct quantities whose evolutionary interpretation is less clear. Unlike many modern methods for re-defining distances, such as UniFrac distances [29] or phylogenetically-defined inner products [38], phylofactorization is principally about discovering phylogenetically-interpretable directions - vectors which characterize primary axes of variation in the community and represented through the contrast basis, a multilevel-factor developed from stepwise selection of factor contrasts, or a basis made of aggregations of the binned phylogenetic units.

### Phylofactorization as a species concept

There is great debate about what constitutes a species in microbes, let alone all organisms. There is a need for objectivity and universality in the definition of “species” and other units in ecology and evolution. The biological species concept is complicated by asexual reproduction. Genetic species concepts are limited by the subjectivity of a sequence-similarity cutoff, such as the 97% sequence similarity commonly used in defining operational taxonomic units or OTUs, which is additionally complicated by the fact that functional ecological similarity may not be uniform at a given sequence-similarity cutoff. Ecological species concepts are often useful once researchers have a clear sense of the functional ecological groups, but it is difficult to objectively define what constitutes an important functional ecological group, especially for taxa whose life histories are unknown. Species concepts coarse-grain the diversity of life in a way that connects our coarse-grained units to biological, ecological, and evolutionary theory. To that end, phylofactorization can be seen as defining a species concept.

Species concepts are fundamental to biology as they partition the diversity of life into units between which we define ecological interactions and within which we define evolution and natural selection. At the heart of species concepts are the operations fundamental to phylofactorization: aggregation, contrast, and an objective function. Species are aggregations of finer units of diversity: individual subpopulations of individual organisms and their individual cells and the cells’ individual genes are all aggregated to define a “population”. Aggregation in a species concept defines a clear partition for later “within-species” contrasts (evolution) and “between-species” interactions (competition & ecological interactions among populations or aggregates of species). A species concept must meaningfully contrast the units of diversity - the biological species concept contrasts species based on reproductive isolation, the genetic species concept contrasts species based on genetic disimilarity, and ecological species concepts contrast species based on distinct functional ecological traits. The objective function in phylofactorization is the theoretical placeholder for a researcher’s “meaningful contrast”. The units for aggregation and contrast must be done in light of some objective, such as a common fitness or pattern of relative abundance within units over time, space, across environmental gradients and/or between experimental treatments. A full theoretical consideration of phylofactorization as a species concept, as it relates to evolutionary and ecological theory, is saved for future research. For the time being, we note that phylofactorization partitions diversity and yields notions of a “species” which can be aggregated and contrasted with other “species”.

Phylofactorization is a flexible species concept, a hybrid of the phylogeny-based phylogenetic species concept [34] and the character-based ecological species concept [48]. After *k* iterations of phylofactorization, the phylogeny is partitioned into *k* + 1 bins of species referred to as “binned phylogenetic units” (BPUs). BPUs are aggregations of the phylogeny which, up to a certain level of partitioning, are more similar to one-another with respect to the aggregation, contrast and objective function, than they are to other groups. BPUs are a coarse-grained way to cluster entities into “units” of organization with common behavior with respect to the ecological pattern defined in the objective function. Phylofactorization defines functional groups based on phylogenetic partitions and a similar association with some ecological pattern of interest. Consequently, phylofactorization can be seen as an ecological species concept constrained to a phylogenetic scaffold. Whereas the phylogenetic species concept is character-based and pattern oriented, phylofactorization is pattern-based and phylogenetically-constrained. A textbook example of a phylofactorization-derived species are “land-dwelling tetrapods”, a group which can be obtained objectively through phylofactorization and which defines a scale for aggregating and summarizing the pattern of vertebrate species-abundances across land/ water habitats.

Phylofactorization permits optional fine-graining and coarse-graining of our patterns of diversity. Phylofactorization provides an algorithm for identifying relevant units, and those units may be at different taxonomic or phylogenetic depths but species within those units will have shared evolutionary history and similar associations with the ecological pattern of interest. For microorganisms, for which the biological species concept doesn’t apply, the genetic species concept appears too detached from ecology, and the ecological species concept is unavailable due to lack of life history detail, phylofactorization serves as a way to organize diversity for focused between-species interactions and within-species comparisons.

### R package: phylofactor

An R package is in development and, prior to its stable release to CRAN, publicly available at https://github.com/reptalex/phylofactor. The R package contains detailed help functions and supports flexible definition of two-sample tests (the function `twoSampieFactor`), contrast-basis analyses with the function `PhyloFactor`, and generalized phylofactorization of exponential family random variables with the function `gpf`. Phylofactorization is highly parallelizable, and the R package functions have built-in parallelization. The R package in development also works with phylogenies containing polytomies, allowing researchers to collapse clades with low bootstrap support to make more robust inferences. The output from phylofactorization is a “phylofactor” object containing the contrast basis, the BPUs, and other details allowing one to input the object into various functions which summarize, plot, cross-validate, run null simulations, and parse out the information from phylofactorization. Researchers are invited to contact the corresponding author for assistance with the package and how to produce their own customized phylofactorizations - such feedback will be essential for a user-friendly stable release to CRAN.

Until then, the supplemental information contains the data and scripts used for all analyses done in this manuscript in an effort to accelerate method development in this field.

### “Everything makes sense in light of evolution”

Phylogenetic factorization is a new paradigm for analyzing a large class of biological data. Ecological big-data, as Thomas Dhobzansky noted about biology in general, makes sense “in light of evolution”. Phylofactorization extends a broad category of data analyses - two sample tests, generalized linear modelling, factor analysis and PCA, and analysis of spatial and temporal patterns - to incorporate a natural set of variables and operations defined by the phylogeny. Phylofactorization localizes inferences in big data to particular edges or chains of edges on the phylogeny and, in so doing, accelerates our understanding of the phylogenetic scales underlying ecological patterns of interest. The problem of pattern and scale is central to biology, and phylofactorization uses the pattern to objectively uncover the relevant phylogenetic scales in ecological datasets.

## Acknowledgments

This work is published in loving memory of Diana Nemergut. This research was developed with funding from the Defense Advanced Research Projects Agency (DARPA; D16AP00113).

## Table of mathematical notation

| Term | Description |
| --- | --- |
| $A(.)$ | Aggregation operator |
| $C(., .)$ | Contrast operator |
| $\mathcal{F}(\theta)$ | Distribution parameterized by $\theta$ |
| $F_e$ | F-statistic for edge $e$ |
| $K_t$ | Number of edges considered in iteration $t$ of phylofactorization |
| $N$ | Size of a binomial random variable |
| $Q$ | A group $Q = R \cup S$ aggregated at a current or previous iteration |
| $R, S$ | Two groups contrasted containing $r$ and $s$ species, respectively |
| $U, B, P$ | Meta-data subsets for phylofactorization |
| $\mathcal{T}$ | Phylogenetic tree |
| $B$ | $m \times p$ coefficient matrix |
| $W$ | matrix of component scores corresponding to $V$ |
| $V$ | $m$ matrix of contrast basis elements |
| $X$ | $m \times n$ data matrix used for phylofactorization |
| $Y$ | $K \times n$ matrix of component scores, one for each edge considered |
| $Z$ | $n \times p$ matrix of meta-data used in regression-phylofactorization |
| $a$ | Coefficient in aggregation vector |
| $b, c$ | Coefficients in a contrast vector |
| $e_k$ | Edge $k$ |
| $e^*$ | Winning edge |
| $e_t^*$ | Winning edge at iteration $t$ |
| $f(.)$ | Transformation in generalized $f$-mean |
| $g$ | Factor containing two levels, $\{R, S\}$ |
| $i, j, k, l$ | Indexes. Often, $i$ is the index for species and $j$ for samples. |
| $m$ | Number of species |
| $n$ | Number of samples |
| $p$ | Number of meta-data types for each sample |

| Terms | Description |
| --- | --- |
| $q$ | Number of pure aggregates in a basis for $\mathbb{R}^m$ |
| $r, s$ | Numbers of species in groups $R, S$ respectively |
| $s(\cdot)$ | Smoothing spline notation for term in generalized additive model |
| $t$ | Iteration of phylofactorization |
| $x_{i,j}$ | The $i, j$th element of data matrix $\mathbf{X}$ |
| $x_{R,j}$ | Aggregate, $A(\mathbf{x}_j)$ of group $R$ for sample $j$. If $j$ is missing then sample is arbitrary. |
| $x_{S,j}$ | See $x_{R,j}$ above. |
| $x_i$ | A random variable (assumed to be a single species $i$ for arbitrary sample) |
| $[x]_{i,j}$ | $i, j$th entry of data matrix, $\mathbf{X}$ |
| $z_i$ | Column of meta-data matrix, $\mathbf{Z}$ |
| $\mathbf{v}_{Q,i}$ | $i$th element of aggregation basis for set $Q$ |
| $\mathbf{v}_{C_{R\vert S}}$ | Contrast vector splitting groups $R$ and $S$ |
| $\mathbf{v}_{C_e}$ | Contrast vector for edge $e$ (which splits sub-tree into two disjoint groups) |
| $\mathbf{x}_{R,j}$ | $r$-vector containing only the species in group $R$ for sample $j$ |
| $\mathbf{x}_{S,j}$ | See $\mathbf{x}_{R,j}$ above. |
| $\mathbf{x}$ | $m$-vector of species' data for an arbitrary sample |
| $\bar{\mathbf{x}}$ | Sample mean of vector $\mathbf{x}$ |
| $\mathbf{y}_e$ | $n$-vector of component scores for edge $e$ |
| $\mathbf{z}_k$ | Vector of meta-data of type $k$. |
| $\beta_i$ | Coefficients for linear model |
| $\eta$ | Natural parameter for exponential-family random variable |
| $\kappa$ | Scale parameter for Gamma distribution |
| $\pi$ | Number of failures parameter for Negative Binomial distribution. |
| $\rho$ | Probability of success for Bernoulli, Binomial, Negative Binomial distributions |
| $\sigma$ | Standard deviation for Gaussian random variable |
| $\theta$ | Arbitrary parameters for probability distribution |

